# Intramolecular interactions between folded and disordered regions shape ubiquilin structure and function

**DOI:** 10.64898/2026.03.13.711692

**Authors:** Jessica K. Niblo, Nirbhik Acharya, Maxwell B. Watkins, Carlos A. Castañeda, Shahar Sukenik

## Abstract

Multidomain proteins consist of folded domains connected by intrinsically disordered regions. The flexibility afforded by the disordered regions coupled to the structure and surface chemistry of folded regions allows for unique structural and functional features in these proteins. Yet how intramolecular interactions between disordered regions and folded domains affect multidomain protein structure and function remain poorly understood. Here we use a range of biophysical and computational approaches to measure the intramolecular interactions between the folded domains and disordered regions of ubiquilins (UBQLNs) - essential components of protein quality control that shuttle poly-ubiquitinated client proteins to proteasomal degradation or autophagy. Starting with the yeast UBQLN homolog Dsk2, we find that interactions between two folded domains located at the opposite ends of UBQLN bring about a closed conformation. The prevalence of this closed conformation, however, is modulated by intramolecular interactions involving the disordered regions and folded STI1 domain at the center of the protein. Simulations and analysis of UBQLN homologs across multiple eukaryotic lineages reveals that these disordered:folded domain interactions exist in some UBQLN homologs but are absent in others, indicating possible fundamental differences in function among proteins with the same multidomain architecture.

## 1. Introduction

Proteins carry out the vast majority of cellular functions. For well-folded proteins, the structure-function relationship is well established; sequence encodes structure and structure dictates function. Intrinsically disordered protein regions (IDRs), by contrast, lack a stable tertiary structure and instead exist as an ensemble of rapidly converting conformations.^1^ Despite this structural heterogeneity, IDRs play central roles in a range of critical cellular functions including cell signalling,^2,3^ transcriptional regulation,^4–6^ and maintaining homeostasis.^1,7,8^ An emerging paradigm links IDR conformational ensemble directly to function, suggesting that changes to the ensemble can modulate protein activity in a context-dependent manner.^1^

Within the human proteome, only 7% of proteins are completely disordered and exist as a conformational ensemble, while 27% of proteins are considered well-folded with a stable tertiary structure (**Fig. 1A**). The remaining 66% of proteins represent mixed, multidomain proteins, which contain well-folded domains and IDRs within the same polypeptide chain (**Fig. 1A**). Multidomain proteins take on a range of architectures (**Fig. S1**), with 62% containing a single folded domain with one or two terminal IDR(s), 27% contain two folded domains connected by IDRs, and 11% contain three or more folded domains interspaced by IDRs (**Fig. 1B**). The length of these IDRs, which either act as flexible tails or linkers, vary considerably (**Fig. 1C**).

**Figure 1.**
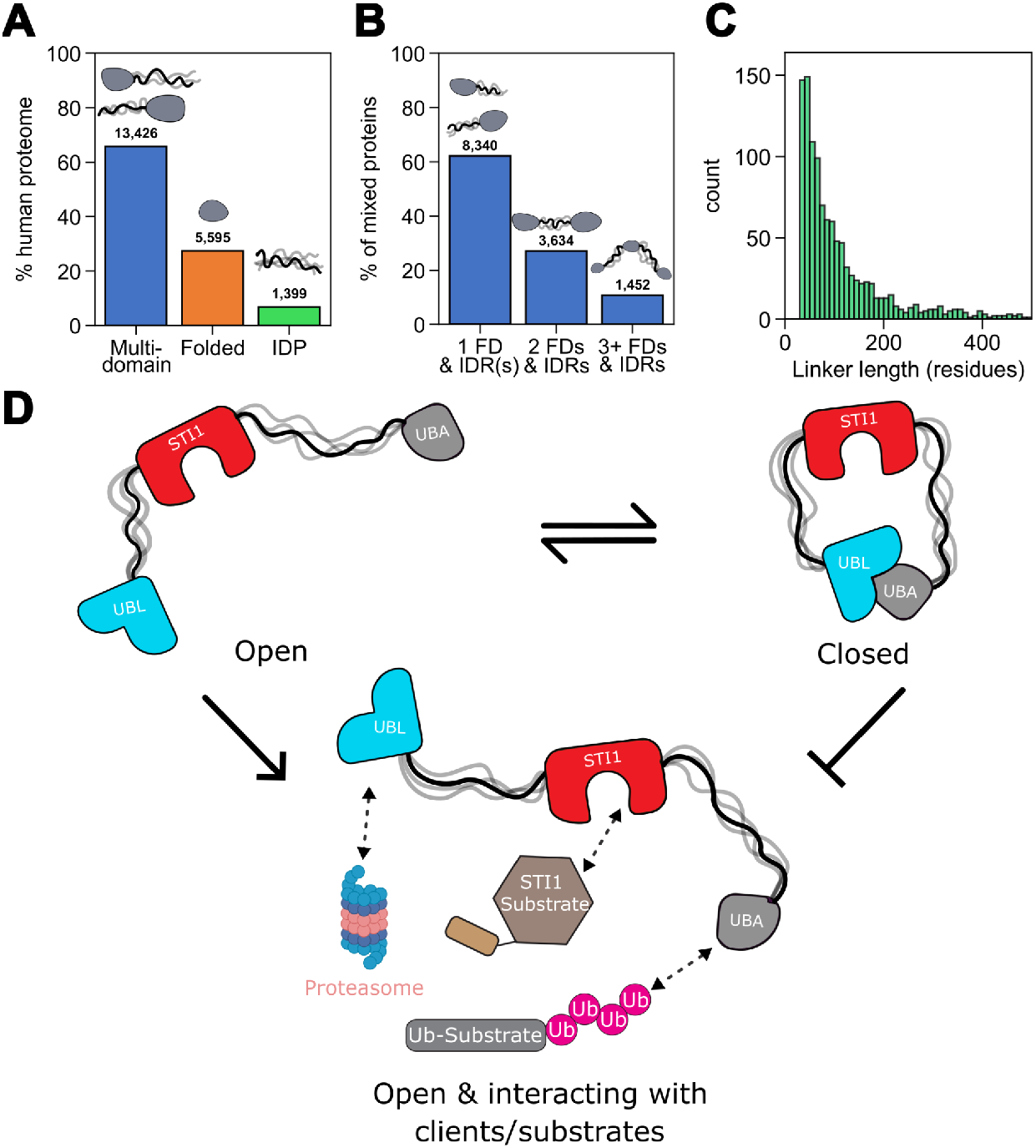
Multidomain proteins are prevalent in the human proteome. (**A**) Breakdown of 20,420 proteins in the human proteome into fully disordered, fully folded, and multidomain architectures. (**B**) Breakdown of the number of folded domains (FDs) in multidomain proteins. Disordered sequences longer than 30 residues were considered IDRs. (**C**) Distribution of the length of IDRs that act as tethers between two folded domains. (**D**) Dsk2, a UBQLN ortholog found in yeast, can exist in either the open or closed (UBL:UBA bound) conformation. Being in an open conformation may allow the protein to functionally interact with ubiquitinated proteins and proteasomes.

In multidomain proteins, the disordered linkers connecting folded domains have competing effects on intramolecular interactions. For one, by tethering two domains together, an IDR enforces a high effective concentration (C_eff_) that can drive intramolecular binding even when binding affinity is modest.^9^ Conversely, the disordered linker can interact with its tethered folding domains, competing with domain-domain contacts. Additionally, IDRs are known to host “stickers” - short linear motifs^10–14^ or folded domains^15,16^ that can bind other proteins to induce a network of self- or hetero-assembled condensates.^17,18^ Together, these inter- and intramolecular interactions can compete with each other to introduce functional complexity: When folded domains are engaged in intramolecular contacts, their ability to bind external partners is reduced, while intermolecular interactions disrupt intramolecular interactions. This inherent competition determines the functional state of the protein, with shifts in the conformational equilibrium controlling domain availability.

Because of these competing effects, switching between distinct intramolecular conformations allows multidomain proteins to regulate activity, specificity, and responsiveness to cellular cues.^19,20^ For instance, the chromatin remodeler ISWI transitions between an auto-inhibited closed state where the termini are bound to each other to an active open conformation when its inhibitory AutoN domain is displaced by histone H4 tail binding.^21^ Similarly, in the epigenetic regulator UHRF1, a short disordered linker transiently occupies a binding groove, resulting in closed conformations that support histone recognition, while linker displacement redistributes the conformational ensemble toward open states with reduced histone-binding efficiency.^22^

Ubiquilins (UBQLNs) provide a clear example in which intramolecular interactions can regulate multidomain protein function. UBQLNs play a key role in protein homeostasis by binding ubiquitinated proteins destined for degradation using a C-terminal UBA domain, and shuttle them to the proteasome by binding it through a folded UBL domain located at the N-terminus (**Fig. 1D**). The terminal UBL and UBA domains are known to bind to one another intramolecularly,^23–26^ forming a closed state (**Fig. 1D**). The UBL and UBA domains are connected by disordered regions that are interspersed by one or two folded STI1 domains in the middle of the sequence. While there is limited structural and dynamical information about STI1 domains,^27^ they are known to engage with chaperones^28,29^ and mediate interactions with client proteins.^30^ Using nuclear magnetic resonance (NMR) spectroscopy, small angle X-ray scattering (SAXS), and coarse-grained simulations, we recently demonstrated that intramolecular interactions between IDRs and STI1 exist in the yeast UBQLN Dsk2. This work also suggests that Dsk2 exists as a weighted ensemble composed of open (UBL-UBA unbound) and closed (UBL-UBA bound) states (**Fig. 1D**).^23^ However, the nature of the intramolecular interactions between Dsk2 folded domains (UBL, UBA, and STI1), its IDRs, how they influence the balance between open and closed states, and how this affects function remains unknown.

To understand how intramolecular interactions shape UBQLN structure and function, we first survey the intramolecular interactions of the yeast Dsk2 UBQLN ortholog using a combined experimental and computational approach. We find that Dsk2 primarily exists in a closed conformation stabilized by UBL:UBA and IDR:STI1 interactions, and that disrupting this closed conformation enhances binding to ubiquitin, a UBA binding partner. Systematic deletion of IDR segments shifts the conformational ensemble of Dsk2 towards open states, revealing a hierarchy of stabilizing interactions. We extend our study to other UBQLN homologs across plants, invertebrates, and vertebrates, revealing that IDR:STI1 interactions are broadly conserved despite sequence divergence. Together, we propose a regulatory mechanism in which binding partners that engage folded domains or IDR motifs shift the conformational equilibrium to regulate UBQLN activity.

## 2. Results

### 2.1 Dsk2 binding is regulated by protein topology

Dsk2 is a yeast UBQLN ortholog and serves as critical ubiquitin-binding shuttle protein^31–33^. Dsk2 (373 amino acids, **Table S1**) contains terminal UBL and UBA domains that bind to proteasome receptors and ubiquitinated substrates, respectively (**Fig. 1D**). When expressed as isolated, untethered domains, UBL and UBA interact with a *K*_d_ of ~ 80 μM, which is weaker than UBA’s affinity for ubiquitin (*K*_d_ = ~ 10 μM)^25,34^. While this untethered binding affinity suggests that UBL:UBA complexes would be weakly populated, tethering these domains dramatically increases their effective concentration, C_eff_,^9,23^ shifting the conformational equilibrium of Dsk2 towards the closed state. We therefore first determined what fraction of the full-length Dsk2 population exists in this closed conformation.

To determine the fraction of population in the closed vs. open topology, we performed small-angle X-ray scattering (SAXS) experiments and correlated these with coarse-grained CALVADOS3 simulations of full length open and closed Dsk2 (**Fig. 2A**). In these simulations, folded domains are held in their correct shape by elastic constraints, while the rest of the protein is treated as disordered. For the open state, when UBL and UBA domains were unbound, the average predicted radius of gyration (R_g_) was 47.0 ± 0.2 Å. For the closed state, additional elastic constraints were applied to UBL and UBA to maintain the domains in the bound orientation observed in the crystal structure of the UBL:UBA complex (PDB: 2BWE).^25^ These closed simulations resulted in an R_g_ of 34.8 ± 0.1 Å. Experimentally, SAXS measurements revealed an R_g_ of 37.2 ± 0.5 Å for the full-length Dsk2 (FL) (**Fig. 2A**, grey), suggesting that Dsk2 exists as an ensemble of open and closed conformations. Reweighting the open and closed simulations to match the experimental R_g_ yielded an optimal distribution of 20% open and 80% closed conformations for Dsk2 FL (**Fig. 2B**).

**Figure 2.**
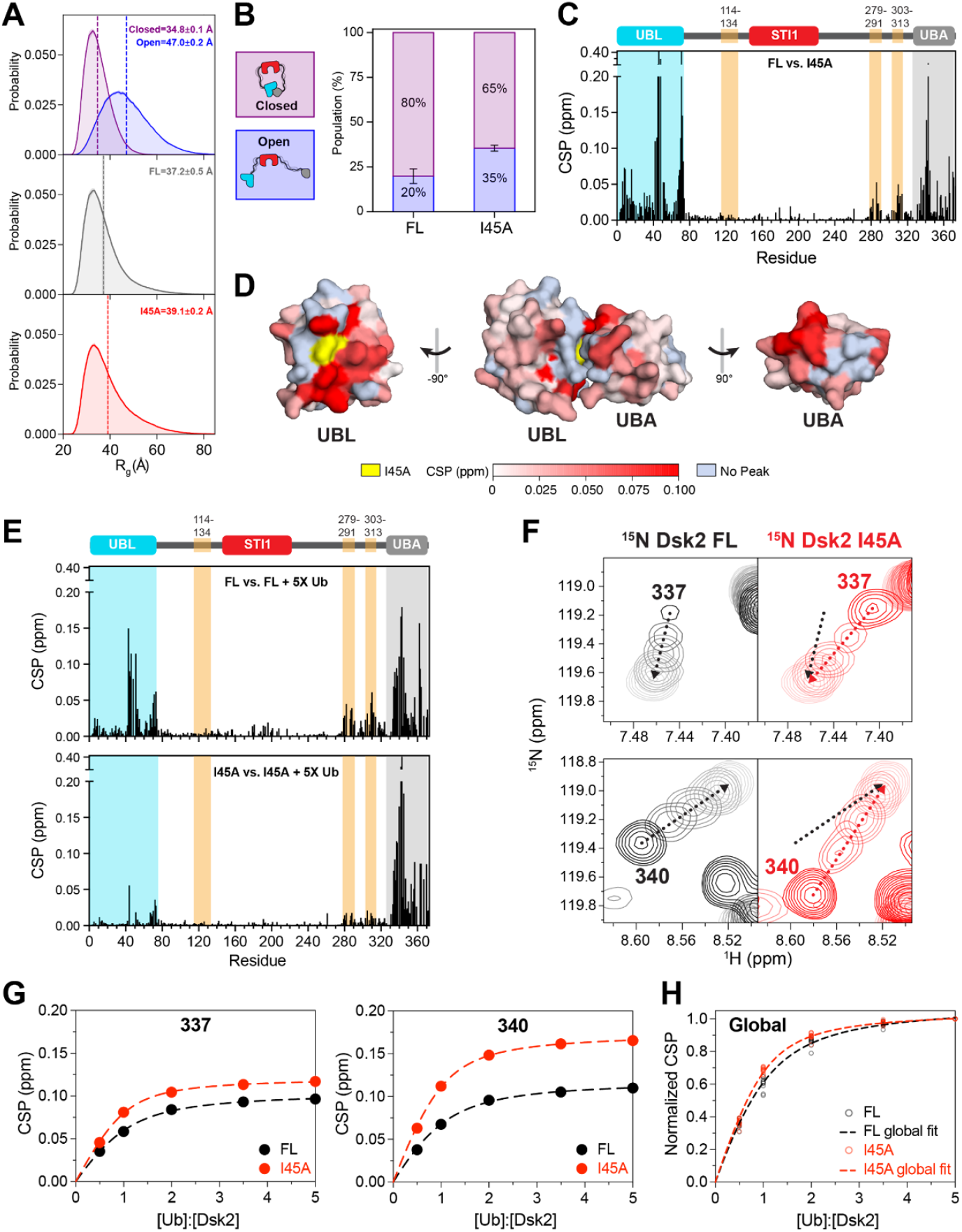
Structural opening of Dsk2 enhances substrate interaction. (**A**) Simulation-derived R_g_ distributions for closed (purple) and open (blue) Dsk2 conformations flank the experimental SAXS-derived R_g_ of the full-length Dsk2 (FL) and the I45A mutant. (**B**) Breakdown of open:closed populations based on simulated ensemble reweighting to experimental R_g_ values. Error bars represent the spread of population estimates from SAXS R_g_ variability. (**C**) Residue-specific amide chemical shift perturbations (CSPs) between Dsk2 FL and I45A mutant. (**D**) Mapping of CSPs from (C) onto the Dsk2 UBL:UBA structure (PDB: 2BWE). Residues colored white-to-red by CSP magnitude; I45A highlighted in yellow; missing residues in pale blue. (**E**) Residue-specific CSPs upon addition of 5X unlabeled ubiquitin (Ub) to 50 µM ^15^N Dsk2 FL (top) or ^15^N Dsk2 I45A (bottom). (**F**) Representative ^1^H-^15^N HSQC spectral overlays showing peak trajectories for two UBA residues (337 and 340) exhibiting large CSPs upon Ub titration into ^15^N Dsk2 FL (black) or ^15^N Dsk2 I45A (red). Arrows indicate peak movement at increasing Ub concentration (dark to light color gradient represents Ub 0:1 to 5:1). (**G**) Representative CSP titration curves for residues 337 and 340 (from panel F) fitted to a 1:1 binding model (see Methods; individual fits in **Fig. S3**). (**H**) Global K_d_ fitting of normalized CSP titration curves for Dsk2 FL (black) and I45A (red). Data points represent normalized CSPs from selected UBA residues (see Methods) and dashed lines show global fits.

To test whether disrupting the UBL:UBA interaction shifts this conformational equilibrium, we mutated I45, a hydrophobic patch residue on the UBL domain that is crucial for both intramolecular UBL:UBA contacts^25^ and binding to proteasome subunits.^26^ The Dsk2 I45A mutation perturbs binding to the proteasome,^33,35^ indicating the importance of this residue at the UBL binding interface. SAXS measurements of I45A resulted in a R_g_ of 39.1 ± 0.2 Å (**Fig. 2A**, red), which is larger than that of the full length protein and closer to the simulated open state. Reweighting the open and closed simulations to match the experimental I45A R_g_ revealed an elevated 35% open population (**Fig. 2B**), consistent with impaired UBL:UBA interactions promoting conformational opening. To validate this, we collected ^1^H-^15^N HSQC spectra of Dsk2 FL and I45A mutant and mapped residue-specific chemical shift perturbations (CSPs) across the protein (**Fig. 2C, S2**). Residues exhibiting the largest CSPs clustered predominantly within the UBL and UBA domains. Mapping these CSPs onto the UBL:UBA crystal structure confirmed that high-CSP residues localize to the UBL:UBA binding interface (**Fig. 2D**), confirming that the I45A variant disrupts UBL:UBA interactions. Additionally, several UBA domain resonances for the I45A variant were shifted toward their positions in the isolated UBA spectrum (**Fig. S2**), consistent with disruption of the intramolecular UBL:UBA interaction and a shift toward more open (UBL:UBA unbound) conformations. Although our NMR results indicate that I45A impairs the UBL:UBA interaction, the SAXS-simulation-derived population distribution shows only 35% open conformation for I45A (**Fig. 2B**). While this represents a twofold increase relative to FL, it remains substantially below complete opening, suggesting that additional factors maintain partial closure in the I45A variant. We hypothesize that the intramolecular IDR:STI1 domain interactions, recently characterized in Dsk2,^23^ prevent complete opening of Dsk2 even when the intramolecular UBL:UBA interaction is disrupted. Consistent with this hypothesis, we observed pronounced perturbations in two regions (aa 279-291; aa 303-313) within the IDR (**Fig. 2C**, orange shaded regions).

We next investigated whether the open-closed conformational shift has any functional relevance in Dsk2 binding to protein quality control components. We compared the ability of Dsk2 FL and I45A to interact with the known binding partner of UBA, ubiquitin (Ub). Because Ub engages the UBA domain and not with the UBL I45A substitution, any differences in binding affinity can be attributed to the conformational state of the protein rather than a direct effect of the mutation. Addition of 5X unlabeled Ub to ^15^N Dsk2 FL produced significant CSPs across both the UBL and UBA domains, consistent with Ub displacing the UBL from its intramolecular interaction with the UBA (**Fig. 2E**, top). This displacement also perturbed the same regions in the IDR (aa 279-291; aa 303-313) that were perturbed in the I45A-mediated conformational opening (**Fig. 2C**), suggesting that Ub-mediated conformational opening disrupts additional interactions. In contrast, for ^15^N Dsk2 I45A, where the UBL:UBA interaction is already disrupted, Ub-induced CSPs were largely confined to the UBA domain (**Fig. 2E**, bottom), reflecting a direct Ub:UBA interaction without disrupting other intramolecular interactions.

To quantify changes to binding affinity, we performed NMR titrations of unlabeled Ub into ^15^N Dsk2 FL and ^15^N Dsk2 I45A up to a 5:1 molar ratio (**Fig. 2F; S3A**). We observed that the UBA residues in I45A consistently showed lower *K*_d_ values compared to Dsk2 FL (**Fig. 2G**; **S3B**). Global *K*_d_ fitting of UBA residues (**Fig. 2H**) revealed that the I45A mutant binds Ub with slightly higher affinity (*K*_d_ = 10.3 ± 0.6 µM) than Dsk2 FL (*K*_d_ = 17.0 ± 0.8 µM). While the >1.6-fold difference is modest, it is consistent with a ~2-fold increase in open population in the I45A compared to FL (**Fig. 2B**), suggesting that the closed conformation of Dsk2 FL partially occludes the UBA binding surface. These results demonstrate that conformational opening enhances substrate accessibility and implicate open-closed equilibrium as a potential regulatory mechanism governing UBQLN function.

### 2.2 Dsk2’s STI1 domain intramolecularly interacts with hotspot regions within IDR and modulates UBL:UBA interactions

The IDR regions perturbed upon structural opening (aa 279–291 and 303–313; **Fig. 2C and 2E**) correspond to two of the three hotspots (HSs) within the IDRs of Dsk2 that were recently identified as interaction partners of the STI1 domain (originally referred to as transient helices in Acharya *et al*.^23^). The STI1 domain is a key functional element conserved across all UBQLNs,^36^ appearing as a single copy in yeast Dsk2 and as two copies in other UBQLNs. STI1:HS interactions drive phase separation and Dsk2-mediated proteasome condensation,^23^ but their role in conformational regulation has not been explored. To test whether these interactions modulate the open-closed topology of Dsk2, we deleted the STI1 domain (Dsk2 ΔSTI1) and compared NMR-derived peak intensity ratios against FL across the protein (**Fig. 3A**). As expected, we observed substantial intensity gains in all three HS regions, consistent with loss of STI1:HS interactions. Notably, we also observed considerable intensity gains in the UBL and UBA domains, suggesting that intramolecular UBL:UBA interactions are weakened in the absence of the STI1 domain despite their higher effective concentration due to the shorter total length of ΔSTI1. These observations are consistent with our recent results^23^ that show elevated R_1_ but decreased R_2_ relaxation rates for both UBL and UBA domains when STI1 is removed; such a pattern could mean that the UBL and UBA domains are tumbling in solution as smaller units, i.e., not in as a tightly bound UBL:UBA complex. We therefore propose that STI1:HS interactions stabilize the intramolecular UBL:UBA complex and promote the closed conformation. Loss of these interactions shifts the conformational ensemble toward more open states. These observations indicate a role for STI1:HS interactions in conformational regulation distinct from, yet complementary to, their previously described role in phase separation.

**Figure 3.**
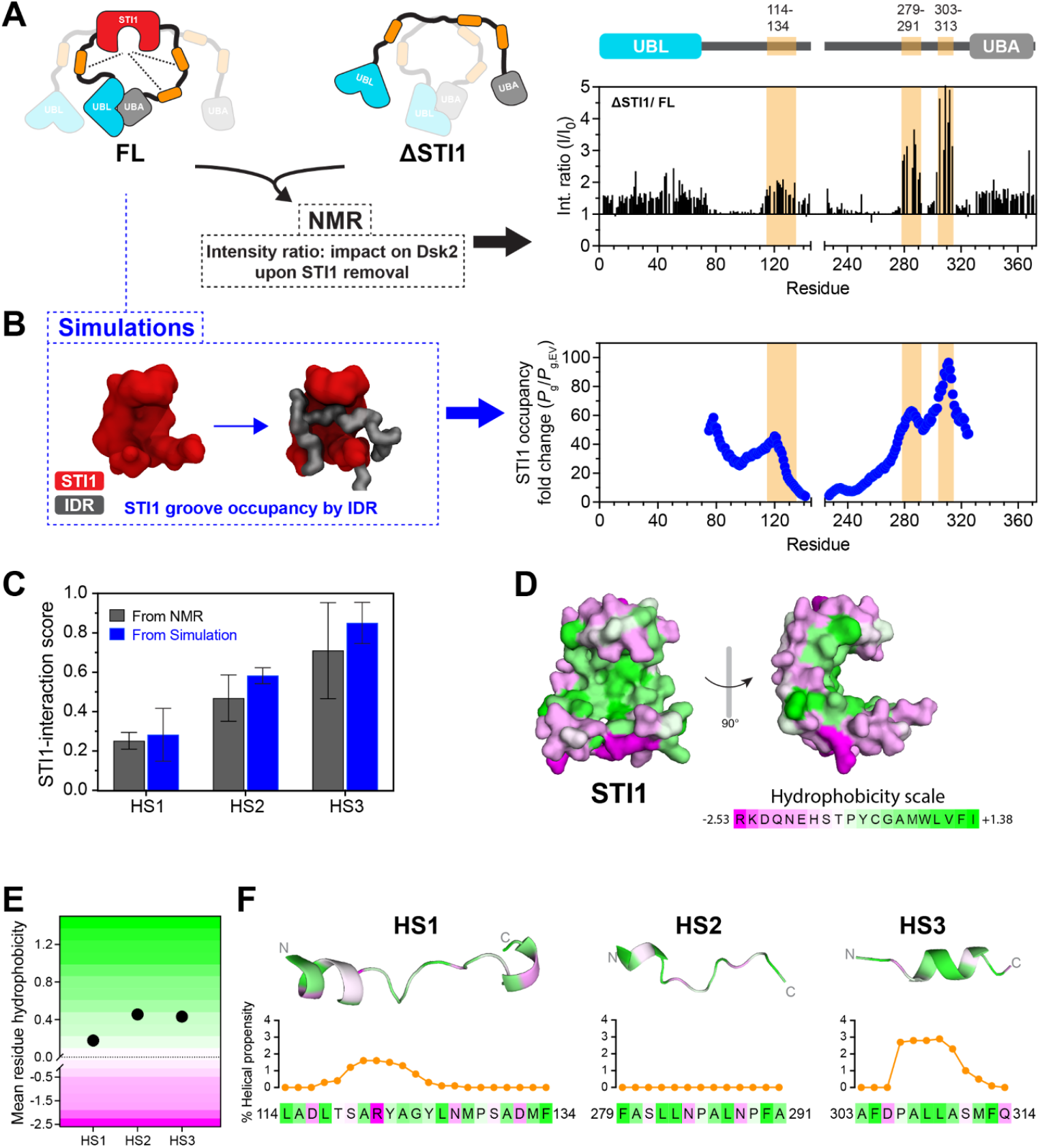
STI1 domain exhibits differential intramolecular interactions with IDR hotspot regions. (**A**) (left) Schematic of Dsk2 FL and ΔSTI1 showing STI1 domain deletion promotes conformational opening (right) residue-specific NMR intensity ratio (ΔSTI1/FL) highlighting intensity gain in IDR regions (orange shading) and UBL-UBA domains upon STI1 deletion. (**B**) (left) Snapshot from simulations illustrating an IDR hotspot occupying the STI1 groove (right) STI1 occupancy probabilities (fold change relative to excluded volume simulations, *P*_g_/*P*_g,EV_) mapping preferential STI1-binding hotspot regions in the IDR (orange shading). (**C**) Normalized STI1-interaction scores for three IDR hotspots (HS1, HS2, HS3) derived from NMR intensity ratios and simulation STI1 occupancy probabilities in panels A and B (see Methods). Data points represent mean ± SD across residues within each HS. (**D**) Surface representation of the STI1 domain of Dsk2 colored according to hydrophobicity scale (AlphaFold P48510), highlighting the hydrophobic groove that engages the IDR hotspots. (**E**) Mean residue hydrophobicity for HS regions show elevated hydrophobicity in HS2 and HS3 compared to HS1. (**F**) Cartoon representations of HS1, HS2, and HS3 of Dsk2 (AlphaFold P48510) colored by hydrophobicity, with residue-specific percent helical propensity shown below in orange (determined using AGADIR^38,39^). Panels D-F use the Eisenberg hydrophobicity scale.^40^

To understand how individual HSs might modulate the open-closed equilibrium of Dsk2 we used simulations to quantify the probability of each IDR residue to occupy the STI1 groove in the open and closed states (**Fig. 3B** schematic). We reweighted the open and closed STI1 occupancy probabilities to match the relative open:closed population ratios determined from the SAXS-derived R_g_ values (**Fig. 2A,B**). It is important to note that in multidomain proteins, intramolecular interactions can occur purely due to positional constraints (*i*.*e*., conformational entropy). We are interested in the effect of sequence chemistry (*i*.*e*., enthalpy) to increase interaction probability beyond this entropic baseline. To measure this excess probability, we performed excluded volume (EV) simulations where all non-bonded interactions were turned off, leaving only hard-sphere repulsions. Using the weighted STI1 groove occupancy probabilities from the full attraction (*P*_g_) and baseline occupancy from the open EV simulation (*P*_g,EV_), we computed the occupancy fold change (*P*_g_/*P*_g,EV_) as the ratio of occupancy probability in the full forcefield to that of the excluded volume simulations (**Fig. 3B**). We observed that the simulation-derived STI1 occupancy fold change shows sharp increases at the three HS regions, mirroring the pattern observed in the NMR intensity ratios (**Fig. 3A**). To enable direct comparison, we normalized both the NMR intensity ratios and simulated STI1 occupancy fold change, and averaged each over the respective HS regions to obtain a relative ‘STI1-interaction score’ for each HS (**Fig. 3C**; see Methods). The experimentally-derived and simulation-derived STI1-interaction scores show strong correlation and consistently reveal that HS3 interacts most strongly with the STI1 domain, followed by HS2 and then HS1 (**Fig. 3C**).

Structural inspection of the STI1 domain reveals a predicted hydrophobic groove (**Fig. 3D**), and sequence analysis of hotspot regions show that HS2 and HS3 carry higher mean residue hydrophobicity than HS1 (**Fig. 3E**). AGADIR predictions indicate that the helical propensity varies among the three HS regions, with HS3 showing the highest propensity, followed by HS1 and HS2 (**Fig. 3F**), consistent with our previous experimental measurements.^23^ Together, these observations suggest that hydrophobicity is the primary determinant of interaction strength for binding to the hydrophobic groove of the STI1 domain, with helical propensity playing a secondary role, consistent with recent findings on human UBQLN2 STI1 domains.^37^

To assess whether the differential STI1-interaction strengths of the three HSs have a corresponding impact on UBL:UBA interactions, we individually deleted each HS region (ΔHS1; ΔHS2; ΔHS3, **Table S1**) and measured chemical shift intensity ratios relative to Dsk2 FL (**Fig. 4A**). The three HS deletions exerted variable effects on UBL:UBA interactions (**Fig. 4B**). Notably, ΔHS3 imparted the highest intensity gains in the UBL and UBA chemical shifts, followed by ΔHS2, with ΔHS1 showing the least impact. These observations are consistent with our initial observation that structural opening of Dsk2 also impacts HS regions, specifically HS2 and HS3 (**Fig. 2C and 2E**). As HS deletions imparted changes in UBL and UBA intensities, the associated CSPs also reported shifts to UBL and UBA resonances at the UBL:UBA interface (**Fig. S4**). These results support a model in which UBL:UBA and IDR:STI1 interactions are coupled in maintaining the closed conformation, such that disrupting one partially destabilizes the other. SAXS measurements corroborate this trend, revealing progressively higher R_g_ values for ΔHS1, ΔHS2, and ΔHS3 relative to Dsk2 FL (**Fig. 4C, S5, Table S2-3**). These SAXS data provide orthogonal validation of conformational opening.

**Figure 4.**
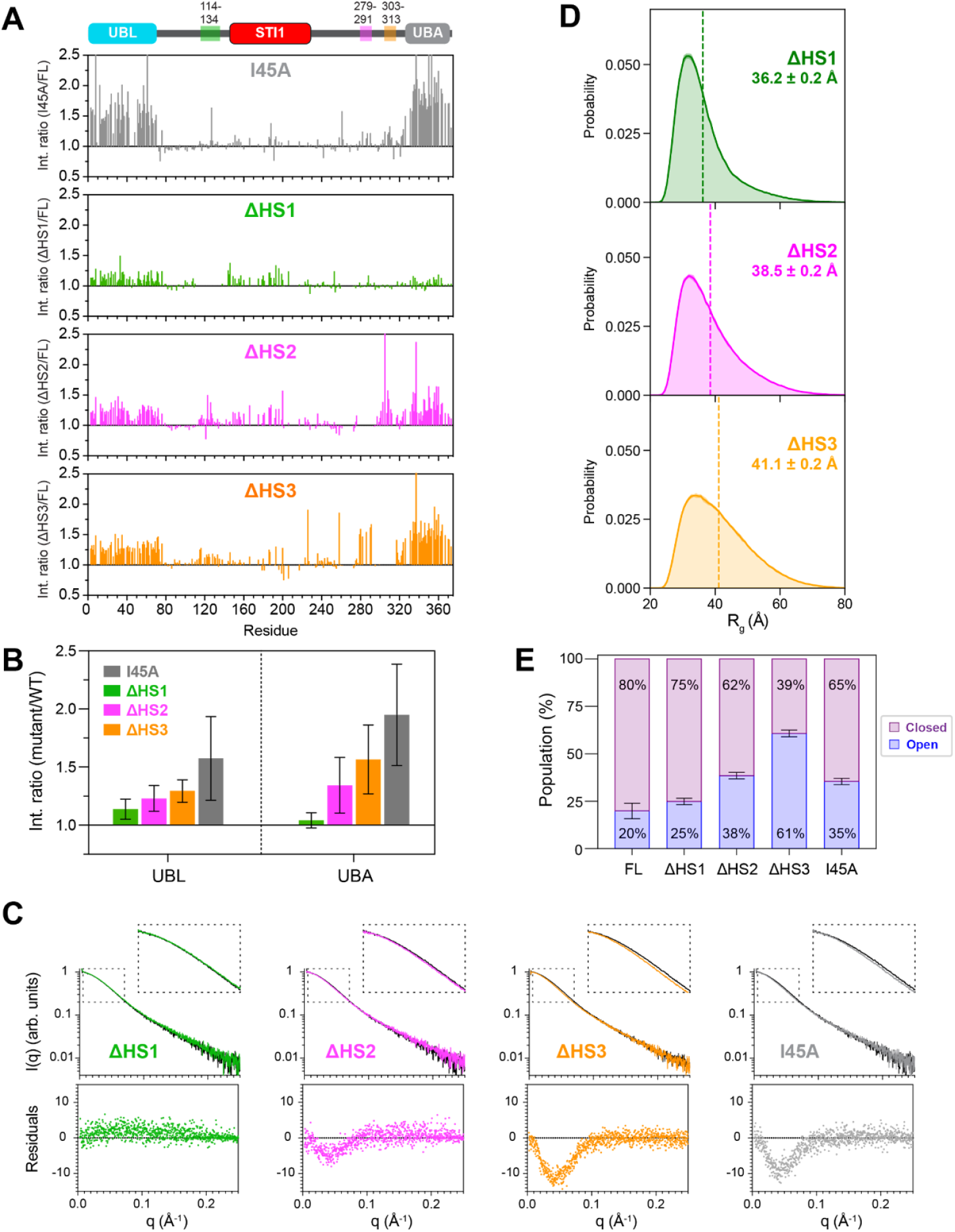
Deletion of hotspot regions in IDR shifts Dsk2 towards open conformation. (**A**) Intensity ratios (I/I_0_) of amide resonances are plotted between Dsk2 FL (I) and variant constructs (I_0_) highlighting dynamic changes in the protein. (**B**) Summary plot showing mean intensity ratio ± SD across residues within UBL and UBA domains from panel A. (**C**) Experimental SAXS scattering profiles for each construct (colored) overlaid with the scattering profile of Dsk2 FL (black) (see Methods). Residuals from profile comparison are shown below each plot. Insets highlight the selected q region (dashed box), where deviations between each construct and Dsk2 FL are most apparent. (**D**) Simulation-derived R_g_ probability distributions reweighted to recapitulate the SAXS-derived R_g_ for ΔHS1, ΔHS2, and ΔHS3. (**E**) Experimental R_g_-based ensemble reweighting compares conformational opening across Dsk2 constructs. Error bars are from SAXS-derived R_g_.

To quantify conformational distributions, we compared experimental, SAXS-derived R_g_ values to simulation-derived values for open and closed states of each HS deletion construct (**Figs. 4D, S6**), as described before (**Fig. 2A**). This analysis revealed a progressive shift toward open conformations for ΔHS1< ΔHS2< ΔHS3 (**Fig. 4E**), consistent with our NMR observations (**Fig. 4B**). These results indicate that the STI1-interaction strength of HS regions is correlated with their impact on intramolecular UBL:UBA interactions. Notably, HS1 is longer in sequence than either HS2 or HS3, yet the HS1 deletion has the smallest effect on conformational opening.

These data suggest that interaction strength, rather than linker length, is the primary determinant of STI1-mediated conformational regulation. Interestingly, a direct comparison between ΔHS3 and I45A variants reveals that ΔHS3 exhibits larger R_g_ and thereby more open conformations (**Figs. 2A and 4D-E**). This is despite the fact that deletion of HS3 shortens the protein and should increase the effective concentration of UBL:UBA interactions. Our data suggest that STI1:HS interactions make a greater contribution to maintaining the compact state than UBL:UBA interactions alone, and may explain why disruption of the UBL:UBA interface via I45A is insufficient to fully open the protein.

### 2.3 Intramolecular IDR:STI1 interactions are conserved across the ubiquilin family

Having established that IDR:STI1 interactions are coupled to regulating the open-closed topology driven by UBL:UBA interaction in Dsk2, we next asked if this mechanism is conserved across the broader UBQLN family. UBQLNs are found across many eukaryotes, and while their core function as Ub-binding shuttle proteins is conserved, the family has diversified in domain architecture. While yeast Dsk2 contains one STI1 domain flanked by two IDRs, most homologs contain two STI1 domains that are connected by three IDRs. This change in protein architecture raises the question of whether the regulatory mechanism observed in yeast Dsk2 is maintained across the UBQLN family, and whether the two STI1 domains interact similarly with the IDRs.

We assembled a set of UBQLN homologs (spanning plants, invertebrates, and vertebrates) and first assessed sequence conservation using sequence alignment^41^ followed by pairwise sequence identity quantification. This analysis revealed that the vertebrate branch shares a moderate identity (>60%) (**Fig. 5A**). Within the vertebrate branch, homologs cluster into their paralogs, with high homology within each group (>85%). Strikingly, invertebrate, plant, and yeast homologs show substantially lower homology relative to vertebrates (~30%), with yeast Dsk2 being the most divergent (**Fig. 5A**).

**Figure 5.**
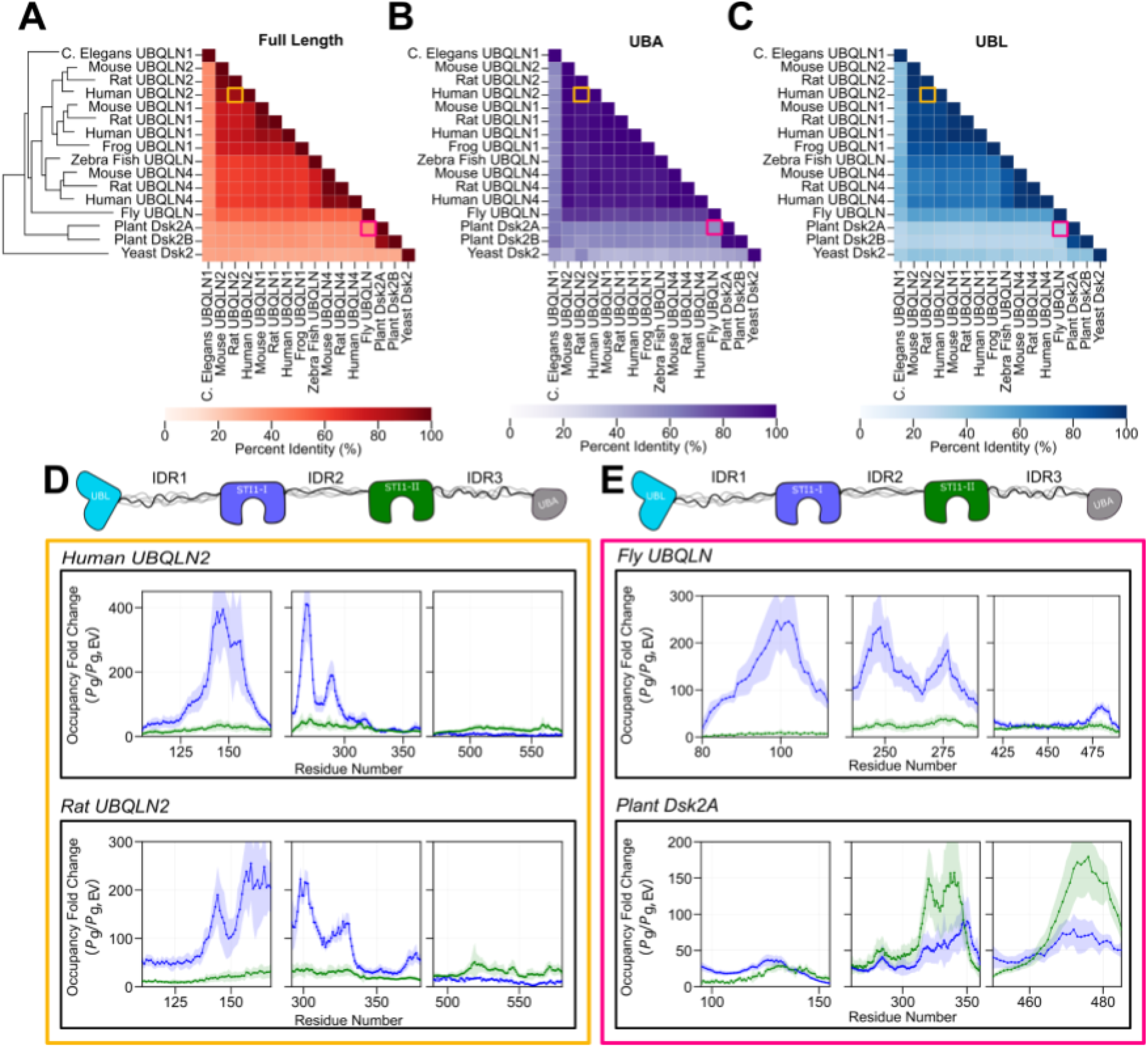
IDR:STI1 interactions are conserved across the UBQLN family proteins. Pairwise sequence comparison of the (**A**) full length protein sequence, (**B**) UBA domain, and (**C**) UBL domain across a range of homologs. Color intensity indicates the percent identity, and homologs are ordered by phylogenetic relationships determined via alignment (tree shown in (**A**)). The occupancy fold change (*P*_g_/*P*_g,EV_) for (**D**,**E**) human UBQLN2 (D, top) and rat UBQLN2 (D, bottom) and fly UBQLN (E, top) and plant Dsk2A (E, bottom). Fold change is shown for both STI1-I (blue) and STI1-II (green) in solid lines, with the shaded bands representing the standard deviation of ten independent simulation replicates.

As UBL:UBA interaction was observed to be one of the drivers to promote closed conformation of Dsk2, we next assessed the conservation of UBL and UBA across UBQLNs. We find the UBA domain has high homology in the vertebrate branch (>80%), and moderate homology extending to invertebrate, plant, and yeast homologs (>50%) (**Fig. 5B**). The UBL domain, while similarly conserved, shows greater sequence variability across the dataset compared to UBA (**Fig. 5C**). Nevertheless, pairwise structure alignments of both the UBL and UBA domains reveal a high degree of structural conservation across all tested UBQLNs (**Fig. S7**). Having established that IDR:STI1 interactions modulate the UBL:UBA interaction in Dsk2, we hypothesized that this regulatory mechanism is maintained across UBQLN homologs.

To determine if the IDR:STI1 intramolecular interactions are conserved despite differences in homology, we perform the same computational analysis that identified Dsk2 hotspots (**Fig. 3B**) on all UBQLN homologies. Remarkably, we find that IDR:STI1 interactions occur in distinct regions of UBQLN IDRs across all homologs. We also point out that the N-terminal STI1-I is the main domain interacting with IDRs, while the C-terminal STI1-II is generally unoccupied (**Fig. S8**).

For orthologs with high sequence homology (**Fig. 5A**), such as human and rat UBQLN2 (89% homology), we observe similar IDR:STI1-I and IDR:STI1-II interaction patterns. Both human and rat UBQLN2 show prominent IDR1:STI1-I and IDR2:STI1-I interactions, with minimal STI1-II engagement and minimal IDR3 interactions (**Fig. 5D**). This interaction pattern extends to fly UBQLN (**Fig. 5E**, top), where STI1-I peaks persist in IDR1 and IDR2 despite lower sequence identity (fly-human UBQLN2 = 53%, fly-rat UBQLN2 = 52%), suggesting IDR hotspots that form strong interactions with the STI1-I domain are conserved across different species. To understand why these interaction patterns persist, we examined the conservation patterns across all three IDRs. Strikingly, all three IDRs differ from one another in their conservation profiles. IDR2 is most conserved, IDR3 shows strong conservation within the vertebrate class but low homology in distant homologs, and IDR1 is weakly conserved with the highest similarity occurring within paralog groups (**Fig S9**). Notably, yeast Dsk2 IDR2, which exists between STI1 and UBA, shows moderate similarity with the IDR2 that sits between STI1-I and STI1-II. This variable similarity is surprising given the conserved IDR:STI1 interactions across the dataset, raising the question of which IDR features are conserved that may drive these interactions. A notable exception to these intramolecular interaction patterns is plant Dsk2A, in which the STI1-II domain is seen to interact more strongly with the IDRs when compared to STI1-I (**Fig. 5E**). Together, these results demonstrate that while IDR:STI1 interactions are broadly conserved across the UBQLN family, the relative contributions of STI1-I and STI1-II can diverge in distantly related homologs.

Given the difference in IDR:STI1-I and IDR:STI1-II interactions, we examined the difference in STI1 homology. Both STI1-I and STI1-II show high conservation in the vertebrate branch (>70%, **Fig. S10A**), with moderate conservation extending to invertebrates and plants (30%, **Fig. S10B**). Strikingly, when comparing STI1-I to STI1-II within individual proteins, most homologs show low intra-protein similarity (~20%, **Fig. S10C**), indicating that STI1-I and STI1-II have stark sequence differences. However, proteins that are seen to have moderate IDR:STI1-II interactions (human UBQLN1, rat UBQLN1, plant Dsk2A, **Fig. S8**) show higher intra-protein STI1-I:STI1-II similarity (~40%, **Fig. S10C**). This suggests that when STI1-I and STI1-II have a moderate sequence similarity, both domains compete to interact with neighboring IDRs. Conversely, when STI1-I and STI1-II have diverged more substantially, STI1-I maintains strong IDR interactions while STI1-II has minimal IDR interactions. Together with the occupancy fold change analysis, these findings indicate conserved hotspot regions driving IDR:STI1 interactions across UBQLNs, and suggest a shared mechanism of IDR:STI1-mediated conformational regulation in UBQLNs, as shown for Dsk2 (**Figs. 3-4**).

## 3. Discussion

Multidomain proteins combine folded domains and IDRs that, through intramolecular interactions, create complex interaction networks that can shape protein function. Here, we have leveraged a combined experimental and computational approach to comprehensively examine how intramolecular interactions regulate the conformational ensemble of yeast UBQLN Dsk2. Consistent with previous results,^23–26^ we find that the UBL and UBA domains interact with one another (**Fig. 2**). SAXS measurements paired with ensemble reweighing of the open and closed coarse-grained simulations reveal that Dsk2 exists primarily in the closed state (**Fig. 2B**) driven by both UBL:UBA and IDR:STI1 interactions (**Figs. 2-3**). Removing IDR:STI1 interactions (through deletions in IDR hotspots, **Fig. 4**) shifts protein topology towards an open conformation where UBL:UBA are not interacting. We show that this shift from closed to open topologies can directly affect UBQLN function by increasing UBA affinity to ubiquitin (**Fig. 2G,H**). Our results highlight how intramolecular interactions between folded domains and IDRs tune the activity of multidomain proteins. Similar observations have been made for a related shuttle protein, Rad23A, where UBL:UBA interactions have been proposed to be auto-inhibitory for Ub recognition.^24^

Having established that IDR:STI1 interactions are coupled to regulating the open-closed conformational ensemble, we extended this analysis to identify similar IDR hotspots in the broader ubiquilin family, which may indicate a conserved IDR:STI1-mediated regulatory mechanism (**Fig. 5D-E, Fig S8**). Despite moderate homology (**Fig. 5A-C, Fig. S9-10**), simulations predict distinct IDR:STI1 hotspots across all studied UBQLNs. This suggests that intramolecular contacts between the STI1 domain and flanking IDRs may represent a general strategy in which the UBQLN family tunes their conformational ensemble. Strikingly, most homologs show preferential IDR interactions with STI1-I with minimal IDR:STI1-II interactions. The STI1-II domain of UBQLN2 has been previously shown to mediate dimerization.^17,42^ Our analysis (**Figs. 5D,E, S8, S10**) suggests that domain divergence may enable functional differences, such as STI1-II serving primarily as a dimerization domain critical for its self-association, while STI1-I functions as an interaction hub for IDR hotspots (**Fig. 5**) and substrates.^37^

Seeing as IDR:STI1 interactions are conserved across UBQLN family proteins, and that disrupting IDR:STI1 interactions may open the conformational ensemble to other binding partners, we examined if functional interaction motifs were embedded in the hotspots with strong IDR:STI1 interactions. Using the ELM database,^43^ we identified short linear motifs (SLiMs) within the IDRs of the UBQLN family and isolated conserved SLiMs that overlapped with predicted hotspot regions (**Fig. 6A**). We found that IDR1 and IDR3, both of which have weak conservation (**Fig. S9**), have very few motifs that are conserved across the dataset. Of the conserved motifs, we observed a few UBQLNs with LIG_LIR_Nem in IDR1, a motif that mediates binding to Atg8 family proteins, which direct ubiquitylated substrates towards degradation (**Fig. 6A**, orange circles).^43,44^ The limited conservation of motifs in IDR1 and IDR3, paired with their minimal IDR:STI1 interactions, suggests these regions may serve paralog-specific functions rather than the conservation of a regulatory role.

**Figure 6.**
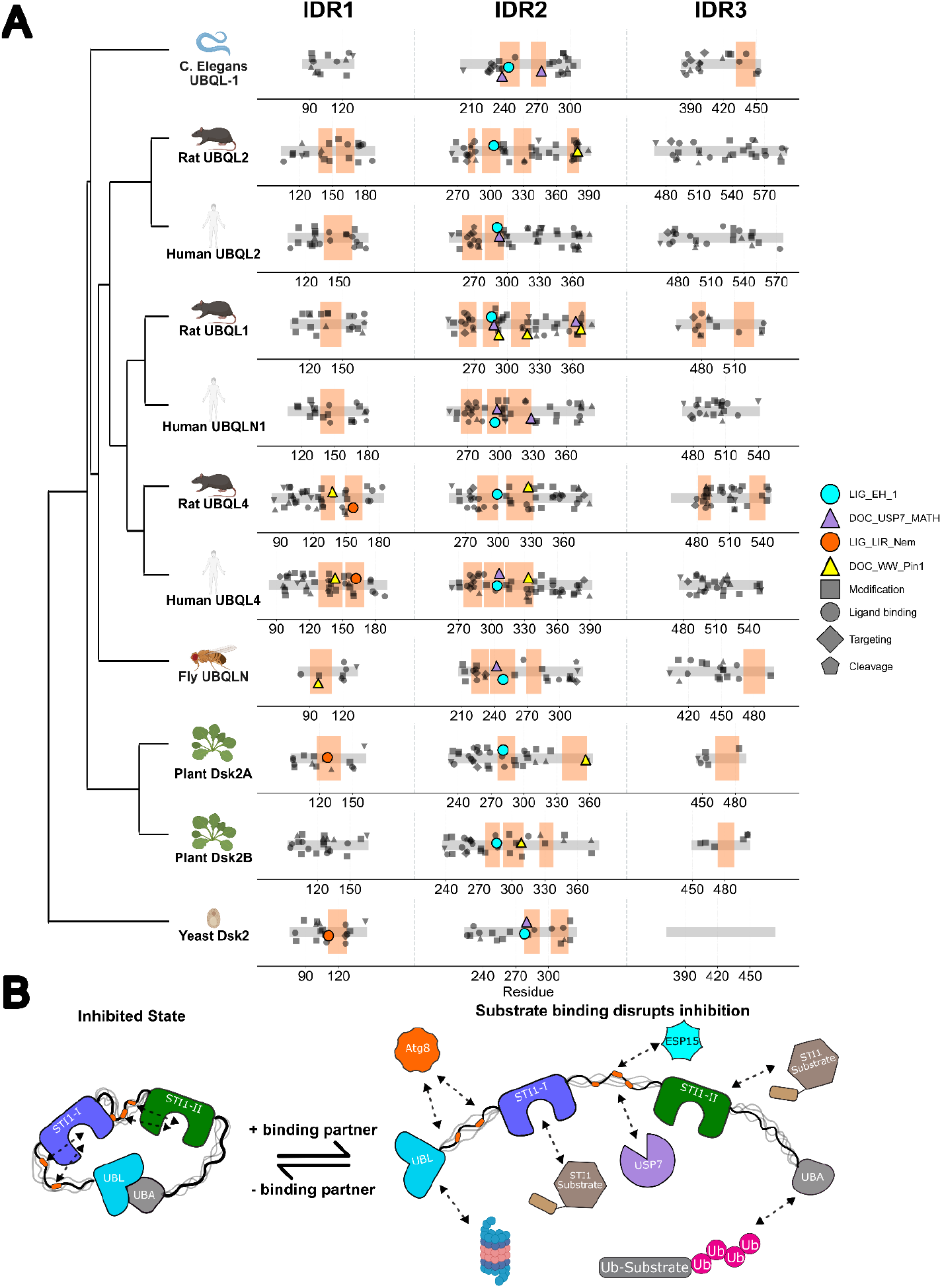
Motifs along UBQLN IDRs. (**A**) Conservation of linear motifs in UBQLN IDR hotspots. Orange boxes indicate hotspot regions within the IDR that have an increased IDR:STI1-1in our simulations (**Fig. S8**). Linear motifs within hotspot regions are shown in large colored markers, and non-conserved motifs outside of the IDR are shown in grey. Marker shape indicates the type of linear motif. (**B**) Schematic showing the proposed interplay between intra- and intermolecular interaction, driven by switching from open to closed topology. Colors of binding partners reflect the color of related motifs in (**A**).

In contrast, we find multiple motifs conserved across IDR2, which has the highest homology and strong IDR:STI1 interactions (**Figs. S8, S9)**. LIG_EH_1, a Asn-Pro-Phe (NPF) motif that binds to the EH-domain of Esp15, is conserved within a STI1-interacting hotspot in IDR2 in all studied homologs. Esp15 is critical to endocytosis and vesicular trafficking.^45^ Interactions between the EH domain and the NPF motif are typically in the mid-micromolar range,^43^ which is comparable to our measured UBA:Ub affinities (**Fig. 2E-H**), suggesting an inherent intermolecular interaction network. Additionally, we find the DOC_USP7_MATH, a motif known to recruit the deubiquitinating enzyme USP7, is conserved in six homologs (yeast Dsk2, fly UBQLN, human UBQLN1, human UBQLN4, rat UBQLN1, and c. elegans UBQLN, **Fig. 6A**). USP7 removes ubiquitin from substrates, reversing the degradation signal, and has been implicated as a key regulator of tumor suppressor pathways.^46,47^ If USP7, Eps15, or other binding partners were to bind to this motif, it would directly compete with IDR:STI1-I interaction hotspots, shifting protein topology towards open conformations, and through this increase or otherwise alter UBA:Ub substrate affinity. Conversely, when the UBA binds a ubiquitinated substrate (**Fig. 1D**), IDR:STI1 interactions weaken, enhancing USP7 accessibility to the motif within the IDR and facilitating deubiquitination of the UBA-bound substrate. Overall, the conservation of these different motifs in STI1-interacting hotspots suggests functional significance that can tune protein activity.

Together, our NMR measurements, simulation data, and the conservation of IDR:STI1 interactions and SLiMs with plausible UBQLN function, we propose a model in which interplay between intramolecular interactions regulate the accessibility of functional motifs to their respective substrates (**Fig. 6B**). The closed state is stabilized by intramolecular UBL:UBA and IDR:STI1 interactions, which further limits the accessibility of these regions to external binding partners (**Fig. 6B**, left). Disruption of either the UBL:UBA interaction, IDR:STI1 interactions, or both through substrate binding (or mutations) promotes conformational opening facilitating interactions with external binding partners involved in substrate processing and delivery. For example, recent evidence suggests that UBQLNs directly interact with non-ubiquitinated substrates via the STI1 domain.^37,48^ Such binding would disrupt IDR:STI1 interactions and also modulate UBL:UBA conformations. Our model provides a framework for understanding how the conformational ensemble of UBQLNs integrates multiple signals to control activity in protein quality control pathways. More broadly, this regulatory mechanism can be a general case for multidomain proteins: tethering creates high effective concentrations favoring intramolecular IDR:folded domain interactions. At the same time, binding partners can shift ensemble dimensions through a concentration-dependent competition for the same interaction sites.

It is important to note some limitations and drawbacks from this study. First, the reliance on coarse grained simulations means that secondary structures, like those present in regions with transient helicity, are not represented in the ensembles. Despite this, we primarily rely on the ability of our simulations to find hotspot regions of IDR:STI1 interactions, which has proven to accurately recapitulate the interacting hotspots detected experimentally by NMR.^23^ We also point out that while we use a two-state model to account for open and closed conformations, there is no evidence of a two-state open/closed equilibrium in Dsk2. Nonetheless, our results showing a shift to more open conformations upon mutations and deletions are not dependent on this equilibrium existing.

## 4. Methods

### Protein Expression and Purification

Yeast Dsk2 constructs and ubiquitin (Ub) were expressed and purified as detailed elsewhere.^23,49,50^ Different domain deletion and mutant constructs of Dsk2 were prepared from the original plasmid using Phusion Site-Directed Mutagenesis Kit (Thermo Scientific) (**SI Table 1**). All Dsk2 constructs were expressed in *E. coli* Rosetta (DE3) cells in Luria-Bertani broth supplemented with 50 mg/L kanamycin and 35 mg/L chloramphenicol grown to OD_600_ of 0.6, induced with 0.5 mM IPTG, and expressed overnight at 18°C for 24 h. NMR active ^15^N labeled protein samples were expressed in M9 minimal media.^17^ Cell pellets were lysed by freeze/thaw in 50 mM sodium phosphate buffer (pH 8.0) containing 300 mM NaCl, 25 mM imidazole, 0.5 mM EDTA, 1 mM PMSF, 1 mM MgCl_2_, and 25 U Pierce universal nuclease. Dsk2 constructs were purified by Ni^2+^ affinity chromatography, followed by 3 h cleavage of the His-SUMO tag with SUMO protease at room temperature during dialysis into 20 mM sodium phosphate (pH 7.2). Cleaved protein was separated from the tag by subtractive Ni^2+^ or Co^2+^ chromatography and further purified by anion exchange. Purified proteins were concentrated using centrifugal concentrators, and concentrations were determined spectroscopically using theoretical extinction coefficients (12950 M^−1^cm^−1^ for Dks2 FL, I45A, ΔSTI1, ΔHS2, ΔHS3; 9970 M^−1^cm^−1^ for ΔHS1). Samples were buffer-exchanged into 20 mM sodium phosphate (pH 6.8) containing 0.5 mM EDTA and 0.02% NaN_3_, then stored at −80°C.

### NMR experiments

NMR experiments were performed at 25°C on a Bruker Avance III 800 MHz spectrometer equipped with a TCI cryoprobe. Samples were prepared in 20 mM sodium phosphate (pH 6.8) containing 0.5 mM EDTA, 0.02% NaN_3_, and 5% D_2_O. Spectra were processed using NMRPipe^51^ on NMRBox^52^ and analyzed with CCPNMR 2.5.2.^53^

### NMR spectra

^1^H-^15^N TROSY-HSQC spectra were acquired with spectral widths of 15 ppm (^1^H) and 27 ppm (^15^N), acquisition times of 200 ms (^1^H) and 46 ms (^15^N), and carrier frequencies of 4.7 ppm (^1^H) and 117.5 ppm (^15^N). Spectra were collected with 16 scans and processed using a Lorentz-to-Gauss window function (15 Hz line sharpening, 20 Hz line broadening) in the ^1^H dimension and a cosine-squared bell function in the ^15^N dimension. Peaks were assigned by transfer from previously deposited backbone assignments (BMRB: 53439; ^23^). Chemical shift perturbations (CSPs) were calculated as Δδ = [(ΔδH)^2^ + (ΔδN/5)^2^]^1/2^, where ΔδH and ΔδN are the ^1^H and ^15^N chemical shift differences, respectively. Peak intensity ratios (I/I_0_) were calculated for a given construct (I) relative to Dsk2 FL (I_0_).

### NMR titration of ubiquitin (Ub) and K_d_ determination

Unlabeled Ub was titrated into 50 µM samples of ^15^N Dsk2 FL or ^15^N Dsk2 I45A, and the binding was monitored by recording ^1^H-^15^N TROSY-HSQC spectra as a function of Ub concentration. CSPs at each titration point were calculated relative to the ligand-free spectrum as described above. Titration data were fit to a 1:1 binding model where the observed CSP (CSP_obs_) is proportional to the fraction of protein in bound state:

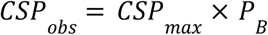

where CSP_max_ is the maximum CSP at saturation and P_B_ is the bound fraction. P_B_ was calculated using the quadratic solution to the 1:1 binding equilibrium^54^:

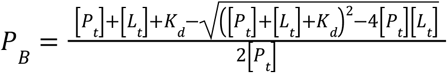

where P_t_ and L_t_ are total ^15^N Dsk2 (FL or I45A) and Ub concentrations, respectively, and K_d_ is the dissociation constant. Individual K_d_ and CSP_max_ values were determined for each residue by nonlinear least-squares minimization using the Levenberg–Marquardt algorithm in LMFIT.^55^ Only UBA residues (327-373) were included in analysis as it is the only binding site for Ub. Fits included only titration points with CSP_obs_ > mean+1 SD (at maximum Ub concentration), with parameter bounds of 0.1–10^6^ µM for K_d_ and CSP_max_ > 0. Residues were excluded if: R^2^ < 0.3, residues with less than three positive CSP points, or fitted CSP_max_ exceeded ten times CSP_obs_. Standard errors were estimated from the covariance matrix.

For global K_d_ determination, UBA residues with CSP_obs_ > mean+1 SD and per-residue R^2^ > 0.3 were fitted simultaneously with a shared global K_d_ while each residue (i) retaining independent CSP_max_:

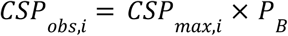

Standard errors for global K_d_ were estimated by bootstrap resampling (100 iterations, resampling with replacement per residue), with the standard error taken as the standard deviation of the resulting K_d_ distribution.

### STI1-interaction score

Relative STI1-interaction scores (**Fig. 3C**) were calculated independently for each hotspot (HS1, HS2, HS3) using two metrics: NMR intensity ratios (I/I_0_ from **Fig. 3A**, where I = ΔSTI1 intensity and I_0_ = FL intensity) and simulated STI1 occupancy fold change (P_*g*_/P_*g,EV*_ from FL simulations in **Fig. 3B**). Each metric was normalized to [0, 1] and averaged across residues within each hotspot, enabling direct comparison between experimental and simulation approaches.

### SAXS data collection and analysis

SAXS was performed at BioCAT (beamline 18ID at the Advanced Photon Source, Chicago) using in-line size exclusion chromatography (SEC) to separate sample from aggregates and other contaminants thus ensuring optimal sample quality and multiangle light scattering (MALS), dynamic light scattering (DLS) and refractive index measurement (RI) for additional biophysical characterization (SEC-MALS-SAXS) (see **Table S3**). The samples were loaded on a Superdex 200 Increase 10/300 Increase column (Cytiva) run by a 1260 Infinity II HPLC (Agilent Technologies) at 0.6 ml/min. The eluent passed sequentially through an Agilent UV detector, MALS detector (18-angle DAWN Helios II, Wyatt), SAXS sample cell, and RI detector (Optilab T-rEX, Wyatt). Scattering was recorded on a Pilatus3 X 1M detector (Dectris) at 3.7 m sample-to-detector distance (q-range 0.0024 or 0.0032–0.33 Å^−1^) with 0.5 s exposures every 1 s during elution (~25 min). Data were reduced in BioXTAS RAW 2.4.0 or 2.4.1 (Hopkins et al., 2017); buffer blanks from regions flanking the elution peak were subtracted from peak exposures to generate I(q) vs. q curves. Peak deconvolution by evolving factor analysis (Meisburger et al., 2016) was performed in BioXTAS RAW. Molecular weights were calculated from MALS/RI data using ASTRA 7 (Wyatt). Data were collected across two beamline sessions (session 1: Dsk2 FL, I45A, ΔHS1, ΔHS2; session 2: Dsk2 FL, ΔHS3). Data within each session were processed identically, and scattering profile comparisons (Fig. 4C) use session-matched Dsk2 FL controls to account for inter-session variability.

### Simulations

We investigated the interactions between intrinsically disordered regions and the STI1 domain for a range of UBQLN proteins by performing single-chain simulations using CALVADOS3_COM_. All interactions are assigned as described in the original CALVADOS framework,^56^ and the elastic network model was independently applied to folded domains as defined in **Table S4** to maintain their folded structure. Using the AlphaFold predicted structures,^59^ full-atom initial conformations were generated using Modeller,^60^ which were then mapped to the CALVADOS3_COM_ coarse-grained representation. Simulations were run at a temperature of 298.15 K, pH of 6.8, and ionic strength of 0.22 M. Yeast Dsk2 was simulated for 70 ns using Langevin dynamics, where the first 3.5 ns was discarded as equilibration and the remaining UBQLN constructs were simulated for 700 ns (**Table S4**). The drag coefficient was set to 0.01 ps^−1^, and the timestep was set to 10 fs. Excluded volume simulations were performed identically, except the non-bonded scaling parameter, λ, and charge, *q*, were each set to 0.

To account for minimal UBA-UBL binding observed in the standard simulations, a second set of simulations was performed where the UBL and UBA domains were placed in a closed conformation. To generate a starting structure, the AlphaFold structure was modified in PyMol such that UBL and UBA were aligned to the crystal structure PDB ID 2BWE.^25^ Following alignment, the structure was minimized using Relax_Amber^61^ and then used to generate ten starting structures using Modeller.^60^ To ensure that UBL and UBA remained in their bound conformation, the elastic bond network was applied to UBL and UBA as a group, and then applied separately to the STI1 domain(s). Simulations were then performed following the same methodology as the open conformation.

All simulation data was obtained from 10 independent trajectories that start from different initial conformations. Each trajectory contained 1,330,000 frames, yielding a total simulation dataset of 13.3 million frames per construct, generated for both full and excluded volume simulations. Ensemble convergence was quantified using the Hellinger distance calculated on the radius of gyration (R_g_) distributions, with a convergence threshold <0.3.^62^

### Simulation Analysis

To quantify the interactions between the STI1 hydrophobic groove and IDRs, we calculate the probability of each IDR residue occupying the STI1 groove, following the approach described in Acharya *et al*..^23^ Each residue of each STI1 domain was either classified as interior (groove-lining) or exterior based on their positions within the coarse-grained structure. The groove center was then calculated as the average position of all interior STI1 residues. For each trajectory frame, we first identified IDR residues that were near both the center of the STI1 groove and at least one interior STI1 residue. Candidate groove-occupying residues were then filtered to remove residues closer to the exterior side of the STI1 domain, as well as those with curvature opposite of the STI1 groove based on PCA of the interior residues. Raw occupancy probabilities were averaged across ten independent simulation replicates and then normalized by the occupancy probabilities obtained from excluded volume simulations to separate sequence-driven interactions from those arising from tethering (**Figs. 3B, 5D,E, S8**). For proteins containing two STI1 domains, occupancy was computed independently for each domain. Hotspots were identified as prominent if they substantially exceeded the surrounding baseline (**Fig. S8**). Full analysis code is available at https://github.com/sukeniklab/UBQLN_interactions_2026.

To estimate the fraction of open and closed conformations, the experimental SAXS R_g_ was fitted as a linear combination of the mean R_g_ value from the open and closed simulations. The resulting optimal weight was then used to construct a weighted ensemble by randomly sampling frames from each simulation in proportion to the optimal weight (**Fig. 2A,B, 4C,D**).

### Bioinformatics Analysis

Disordered regions of proteins of the human proteome were predicted using metapredict v3.0.^63^ Folded domains separated by IDRs shorter than 30 amino acids were considered to be a single domain, and IDRs shorter than 30 amino acids were excluded (**Fig. 1A**). Proteins found to have both IDRs and folded domains were classified as having a mixed architecture. These mixed proteins were then classified into subgroups based on the ordering of their folded and disordered regions (**Fig. 1B**). Patterns were grouped with their reserve equivalents to analyze domain arrangements (**Figure SI1**).

Multiple sequence alignment of UBQLN homologs was performed using MUSCLE.^41^ Pairwise sequence identity was calculated as the percentage of identical residues at conserved positions of aligned sequences. Phylogenetic trees were constructed from these distance matrices using the UPGMA clustering method.^64^ Domain-specific alignments were performed separately for the full-length sequence, UBL, UBA, STI1-I, STI1-II, and IDR regions to assess conservation within domains (**Fig. 5A-C, S10, S9**).

Short linear motifs (SLiMs) were identified using the Eukaryotic Linear Motif database.^43^ Conservation of SLiMs was assessed by determining if motifs overlapped identified STI1-binding hotspots across multiple homologs, with motifs being present in at least three homologs classified as conserved (**Fig. 6A**).

## Data Availability

The datasets and computer code produced in this study are available in the following databases: Simulation analysis and bioinformatics code: Github (https://github.com/sukeniklab/UBQLN_interactions_2026). NMR data (chemical shifts for I45A, HS deletions) are deposited in the BMRB with accession codes TBD.

## Acknowledgements

We acknowledge support from National Institute of Health (NIH) R35 GM137926 to SS, R35 GM158070 to CAC, and postdoctoral support to JKN from the Syracuse University Vice President of Research office. SS acknowledges support from the Alfred P. Sloan Foundation. Support for the Bruker 800 MHz spectrometer with TCI cryoprobe was provided by shared instrumentation NIH Grant 1S10OD012254. We appreciate support from Dr. Charlie Fry at the SUNY-ESF NMR facility. This research also used resources of the Advanced Photon Source, a U.S. DOE Office of Science User Facility operated for the DOE Office of Science by Argonne National Laboratory under Contract No. DE-AC02-06CH11357. BioCAT was supported by grant P30 GM138395 from the National Institute of General Medical Sciences (NIGMS) of the NIH. This study made use of NMRbox: National Center for Biomolecular NMR Data Processing and Analysis, a Biomedical Technology Research Resource (BTRR), which is supported by NIH grant P41GM111135 (NIGMS). We thank Dr. Thuy P. Dao for molecular cloning assistance of tested Dsk2 constructs and scientific feedback. JKN thanks H. Beryl Rappaport for helpful discussions on constructing phylogenetic trees. We thank Syracuse University’s High Performance Computing, including the OrangeGrid cluster (NSF ACI-1341006), for computational resources. We are grateful to Daniel Jeski and Larne Pekowsky (NSF ACI-1541396) for technical support and assistance with cluster usage. The content is solely the responsibility of the authors and does not necessarily reflect the official views of NIGMS or the NIH.

