## Supplementary figures and tables for "Intramolecular interactions between folded and disordered regions shape ubiquilin structure and function"

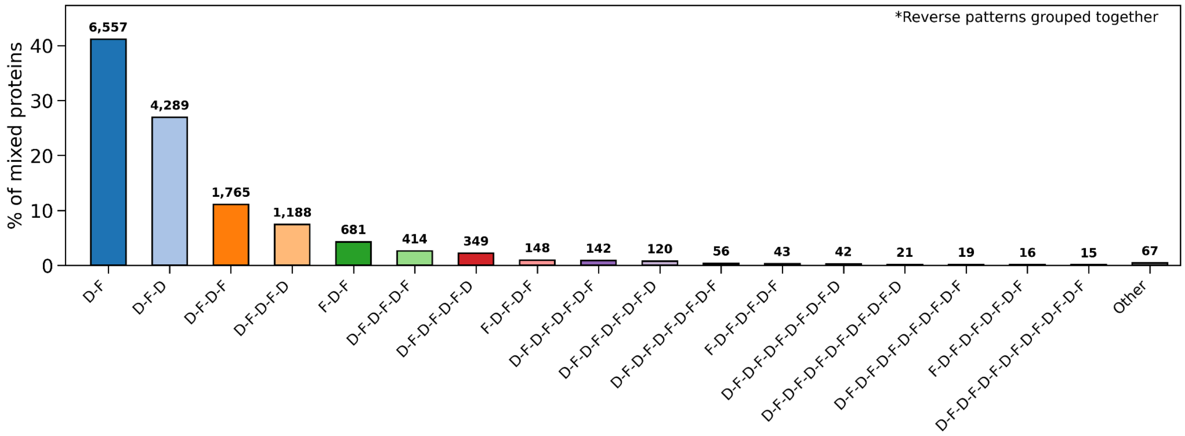


**Figure S1**. **Breakdown of multidomain protein architecture for the 20,420 proteins found in the human proteome.** Disordered and well-folded regions are predicted using Metapredict,[1] and IDRs are taken to be 30 amino acids or longer. The architecture is shown as the x-axis label, with F-D representing a folded domain (F) tethered to a disordered domain (D), with reversed patterns grouped together. Numbers above the bar represent the protein count with the specified architecture. Architectures with fewer than 10 proteins are grouped as other.

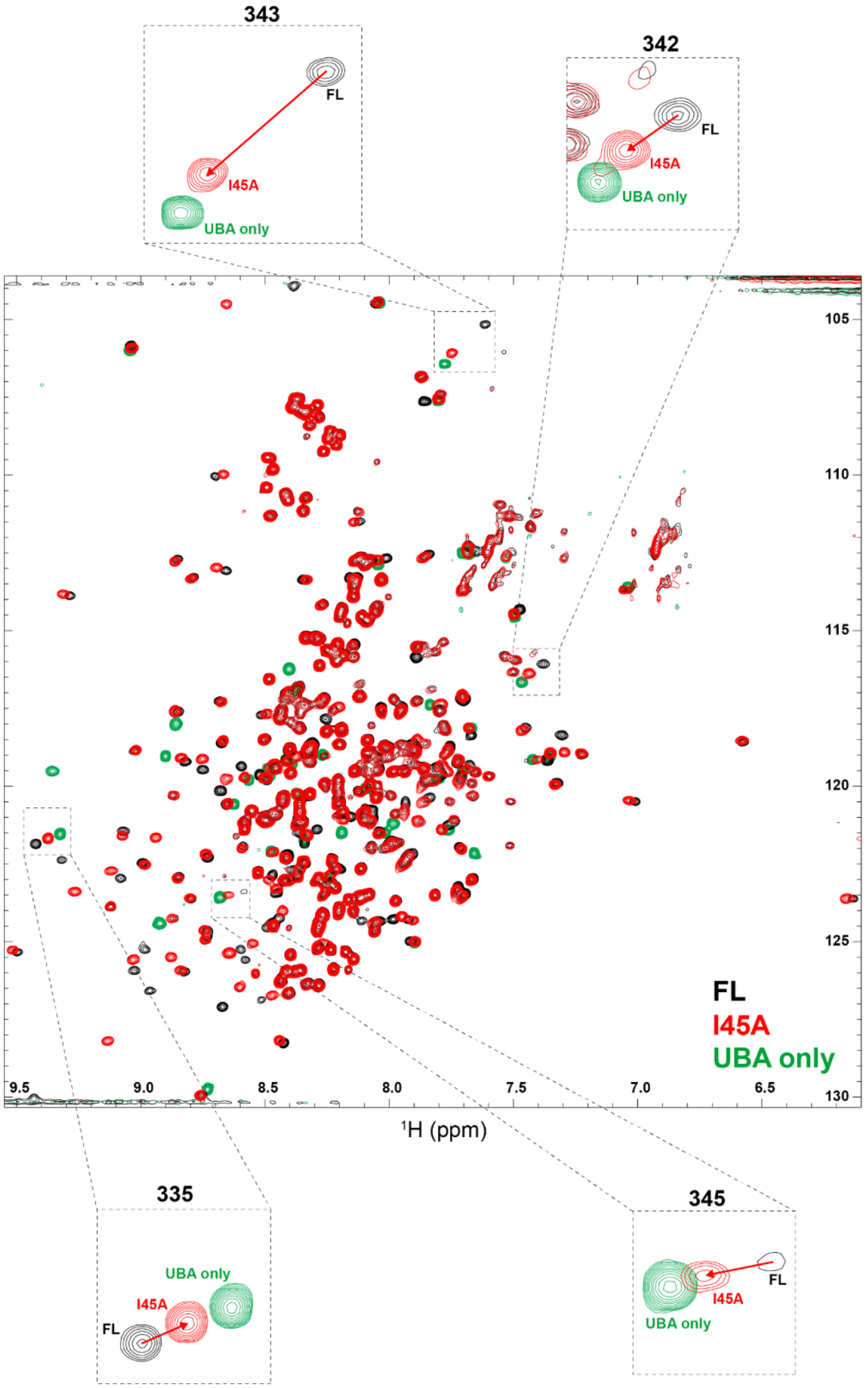


**Figure S2. $^{1}$H-$^{15}$N TROSY-HSQC spectra of Dsk2 FL, I45A and isolated UBA.** Full spectra are compared between Dsk2 FL (black), I45A (red) and isolated UBA domain (green). Zoomed insets highlight selected UBA residues, demonstrating that backbone amide resonances in the I45A mutant shift away from FL positions toward those of the isolated UBA domain, consistent with a more open conformational state. All spectra acquisition parameters (number of scans, receiver gain, etc.) were identical.

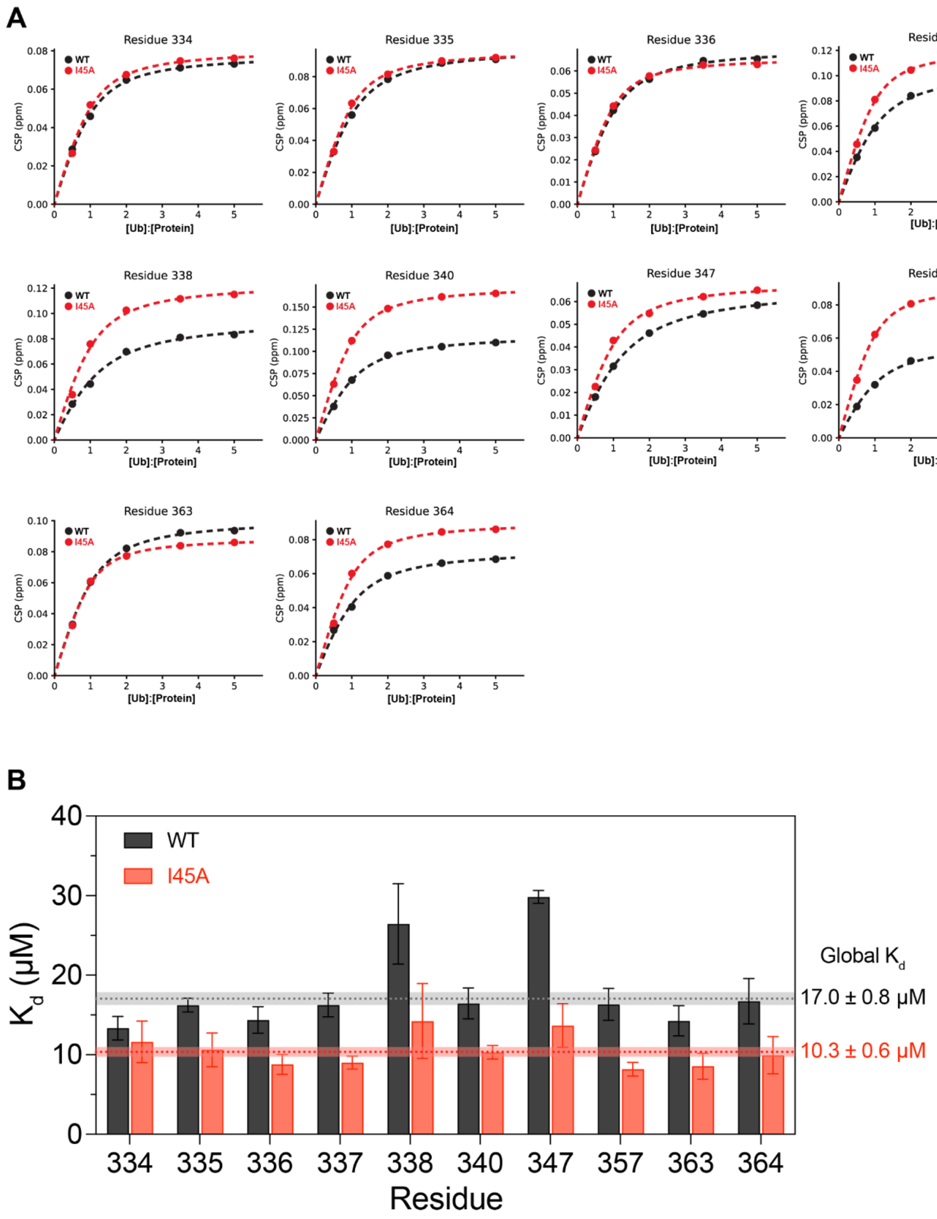


**Figure S3. CSP titration curves and $K_d$ fits for UBA residues in Dsk2 FL and I45A.** (A) CSP titration curves for selected UBA residues in Dsk2 FL (black) and I45A (red) fitted to a 1:1 binding model. Residues were selected based on $CSP_{obs}$ > mean + 1 SD and per-residue $R^2$ > 0.3. (B) Individually fitted $K_d$ values from panel A for Dsk2 FL (black) and I45A (red). Standard errors were estimated from the covariance matrix. Dotted lines indicate global $K_d$ values and shaded regions represent standard errors from bootstrap resampling (see Methods).

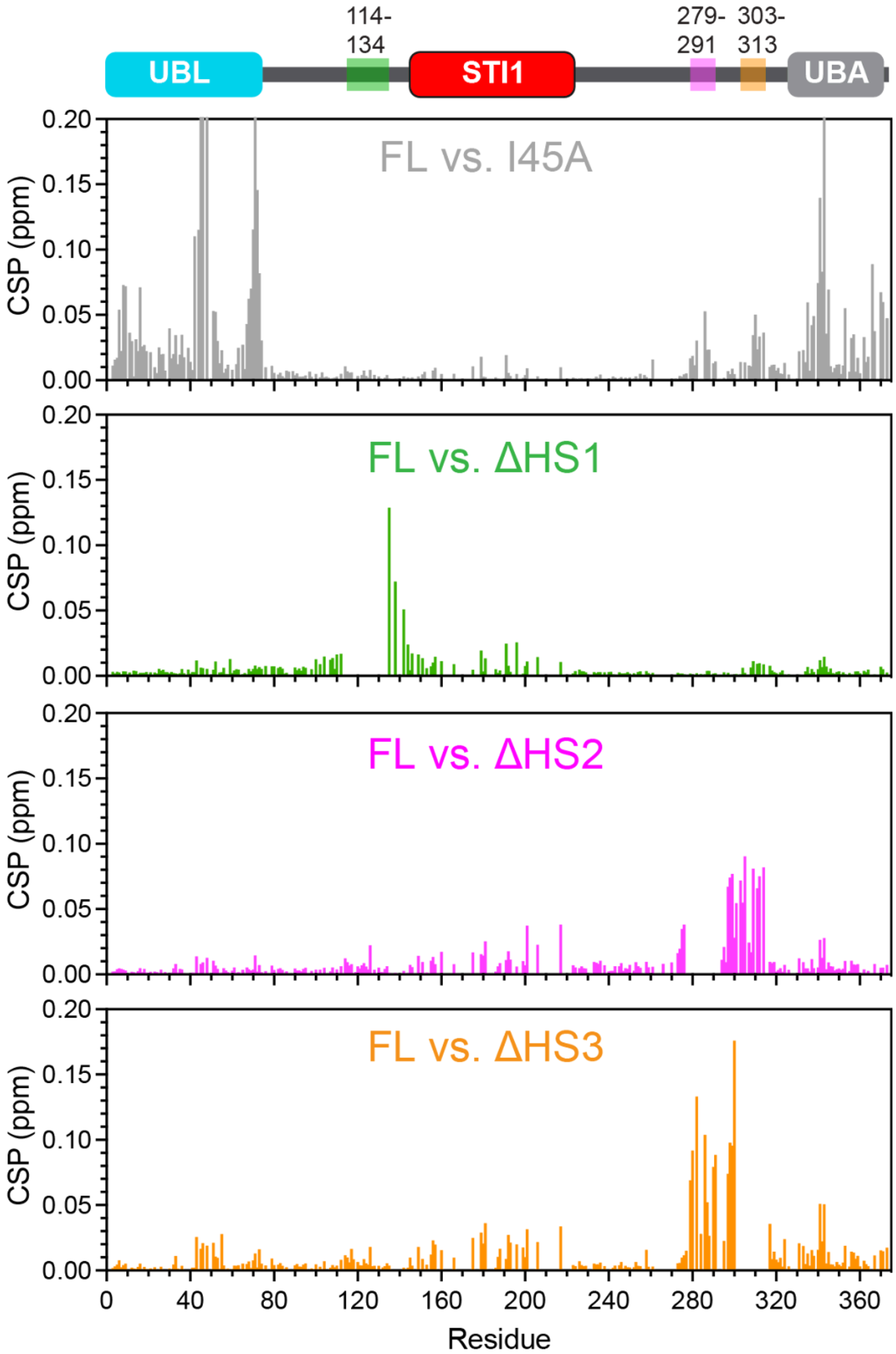


**Figure S4. Chemical shift perturbations for hotspot deletion constructs relative to Dsk2 FL.** Hotspots shown green, pink, and orange rectangles on the domain map at the top of the figure. CSPs for ΔHS1, ΔHS2, and ΔHS3 constructs are compared relative to Dsk2 FL. CSPs for I45A relative to FL (from Fig. 2C) are shown for comparison.

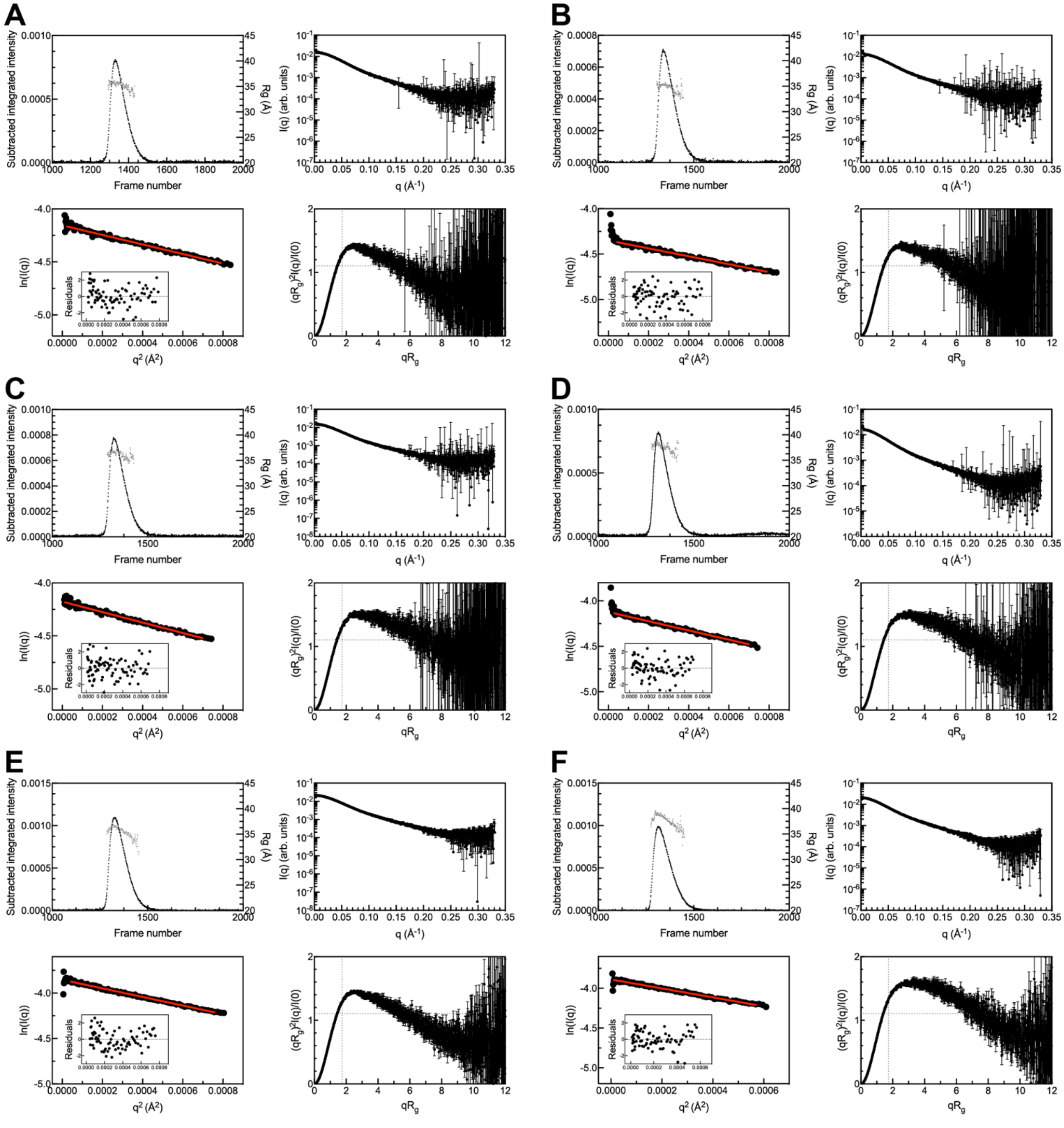


**Figure S5. SAXS data analysis.** SEC-SAXS profiles (top left) for Dsk2 constructs across two beamline sessions (session 1: (A) FL, (B) ΔHS1, (C) ΔHS2, (D) I45A; session 2: (E) Dsk2 FL, and (F) ΔHS3). Buffer subtracted intensity (black) on left y-axis and $R_g$ values (grey) on right y-axis. I(q) vs. q scattering curves (top right) determined from corresponding SEC-SAXS profiles. Red line in Guinier plot (bottom left) is linear fit of ln(I(q)) vs. $q^2$, while inset shows residuals of fit. Dimensionless Kratky plots (bottom right) include dotted lines to indicate where a globular protein would peak.

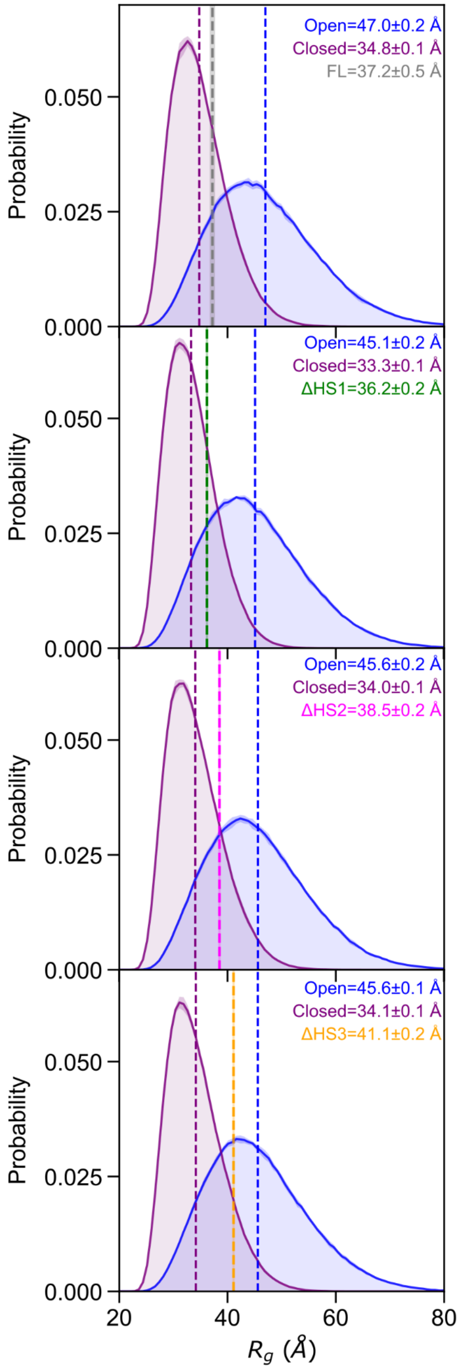


**Figure S6. Simulation-derived $R_g$ distributions for Dsk2 FL and deletion constructs.** $R_g$ probability distributions for the open (blue) and closed (purple) for the Dsk2 FL and deletion variants flank the experimental SAXS-derived $R_g$ value. Average $R_g$ values for open, closed, and SAXS obtained are shown by the dashed lines. Error is shown by the shaded regions. Variants are colored as FL: grey, ΔHS1 green; ΔHS2 magenta; ΔHS3 orange.

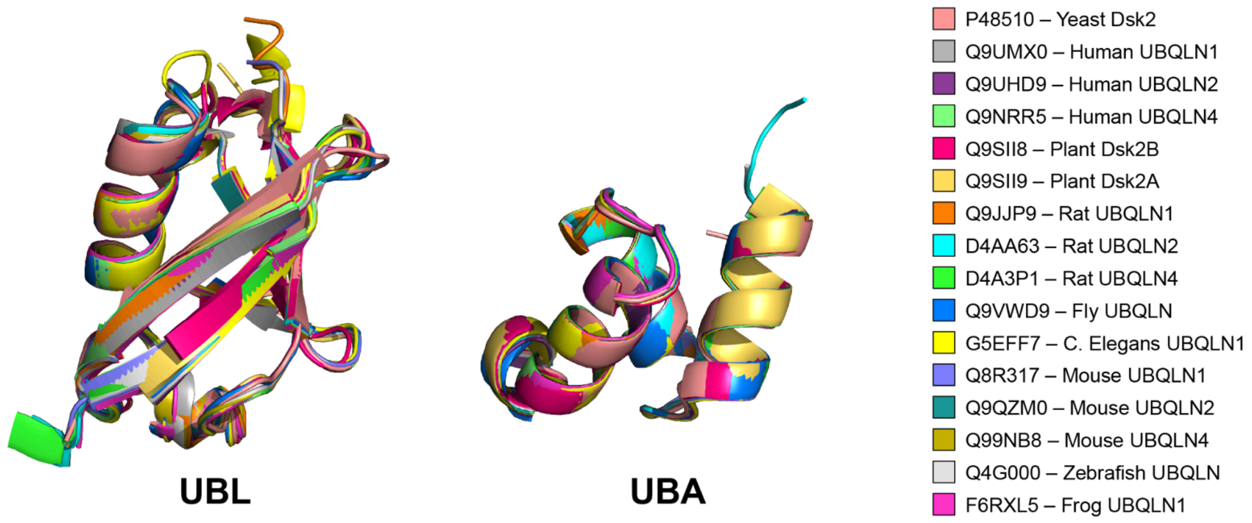


**Figure S7. Pairwise structural alignment of UBL and UBA domains across UBQLNs.** UBL and UBA domain structures from selected UBQLN orthologs (as indicated in the legend: uniprot code- gene name) were superimposed to assess structural conservation. Structures were obtained from AlphaFold and aligned using the TM-align method on Pairwise Structure Alignment tool (https://www.rcsb.org/alignment;[2]). Domain boundaries were set according to **Table S4**. Despite low sequence identity (**Fig. 5B,C**), both domains adopt highly conserved folds across all species examined.

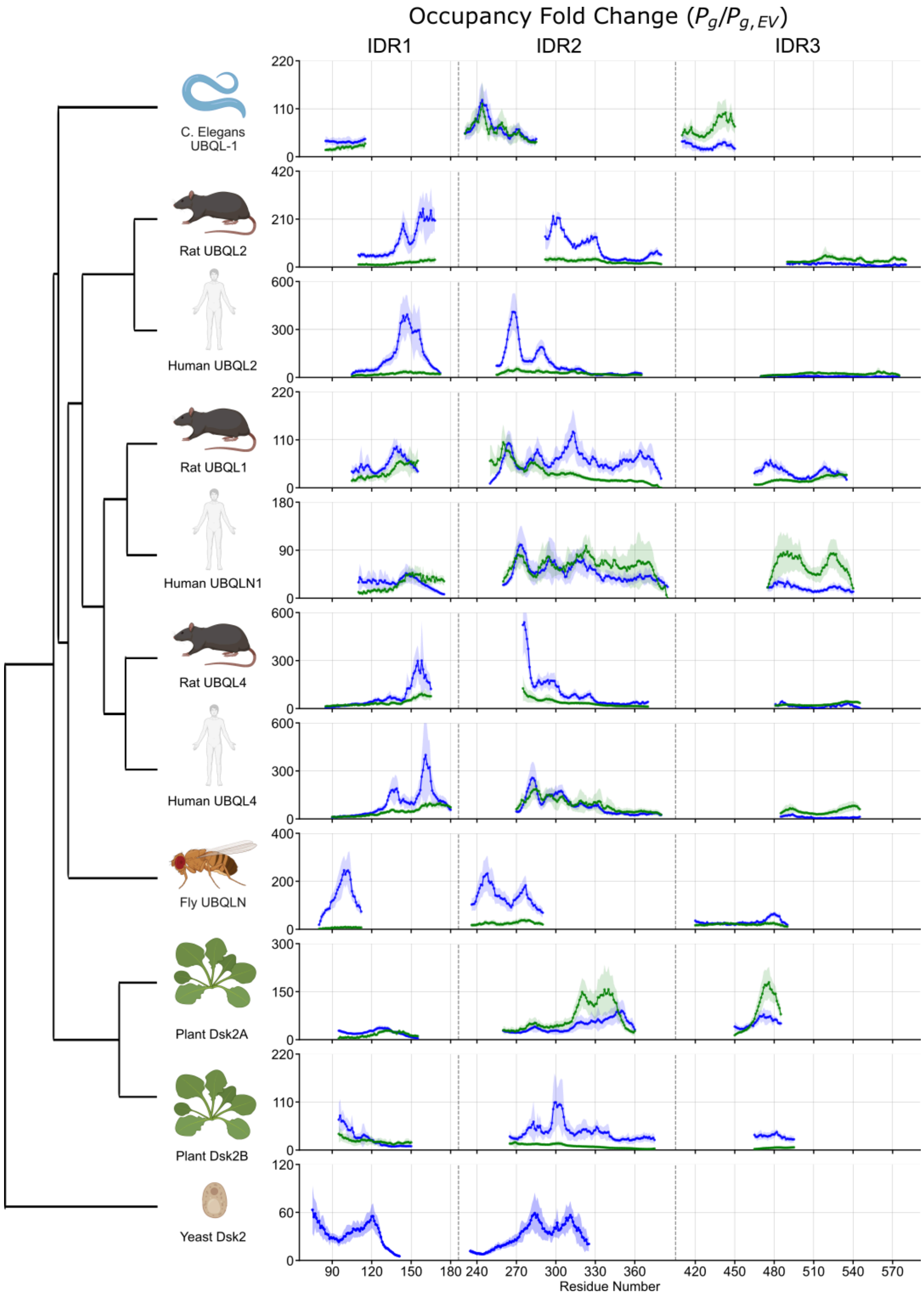


**Figure S8:** STI1 occupancy fold change for the open conformation of several UBQLN proteins. Occupancy probability for each residue of the IDRs to occupy the STI1-I (blue) and STI1-II (green) domain is calculated for the full attractive forcefield and excluded volume forcefield. The occupancy fold change is shown as the ratio of $P_g/P_{g,EV}$ for different UBQLN proteins spanning plants, invertebrates, and vertebrates. Occupancy fold change is shown by solid lines, with the shaded bands representing the standard deviation of ten independent simulation replicates.

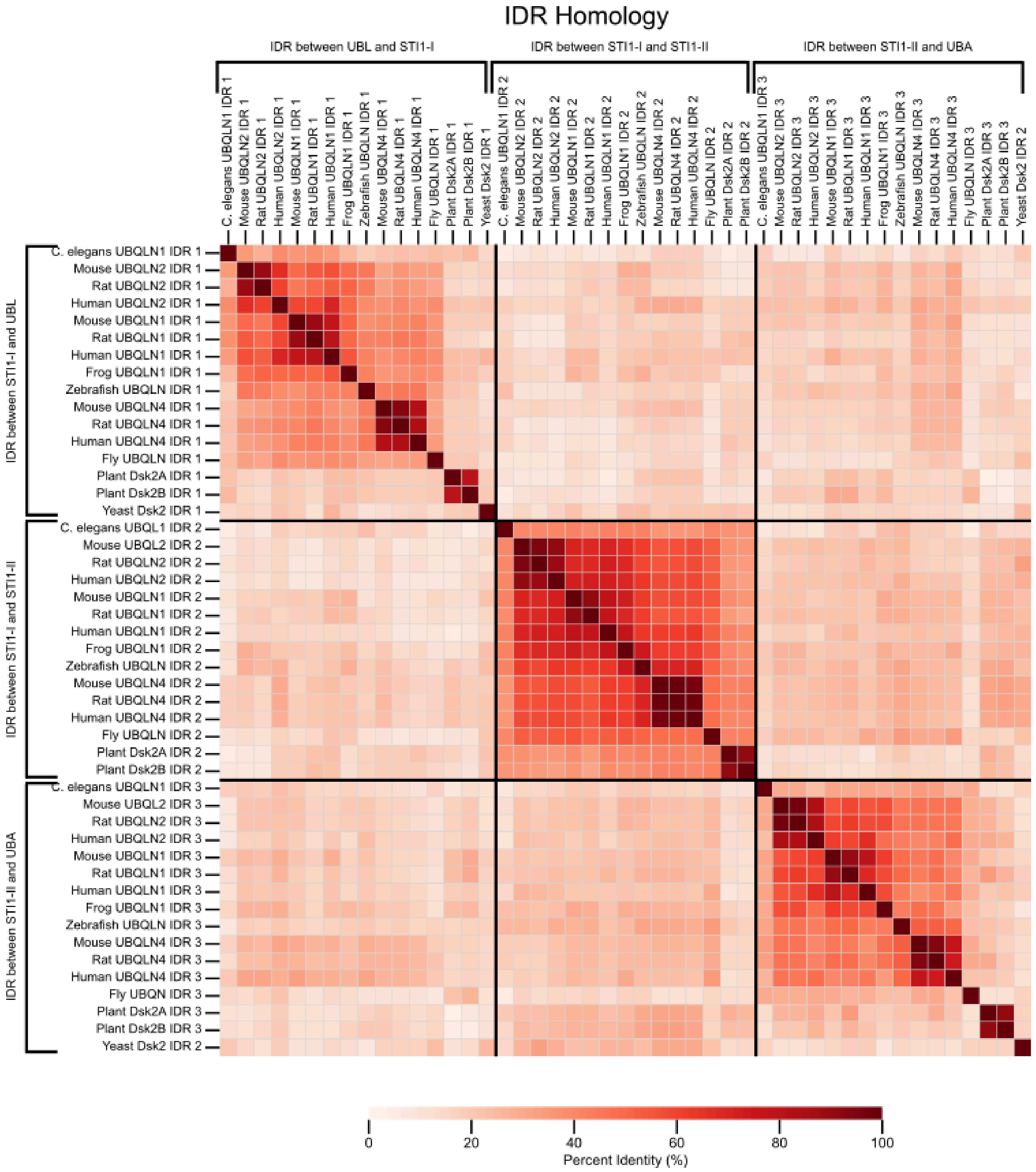


**Figure S9. Pairwise identity of the IDRs of the UBQLN family proteins.** Homology of the IDRs found between UBL and STI1-I (IDR1), STI1-I and STI1-II (IDR2), and STI1-II and UBA (IDR3). Yeast Dsk2's second IDR is positionally equivalent to the IDR between STI1-II and UBA of other organisms (IDR 3). Sequences were aligned using MUSCLE3 and ordered to reflect the phylogenetic relationships established in Fig. 5A of the main text. Percent identity was calculated from pairwise alignments excluding the gap positions.

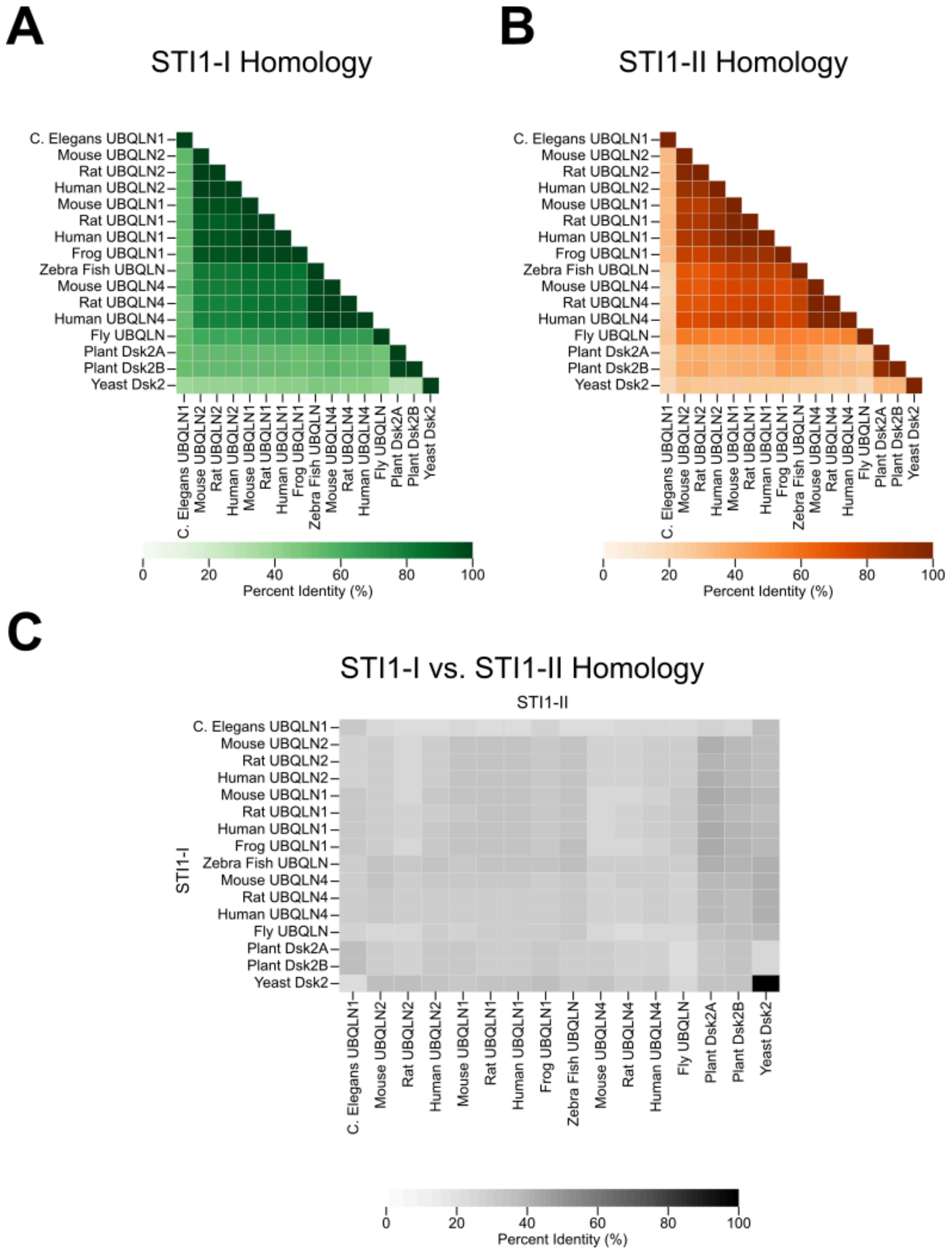


**Figure S10. Pairwise identity of the STI1-I, and STI1-II of the UBQLN family proteins.** Homology of the A) STI1-I, B) STI1-II, and C) Cross-domain comparison of STI1-I vs. STI1-II. As yeast Dsk2 only contains one STI1 domain, it is included in both (A) STI1-I and (B) STI1-II comparisons, and on both axes of the cross comparison. Sequences were aligned using MUSCLE and ordered to reflect the phylogenetic relationships established in Fig. 5A of the main text. Percent identity was calculated from pairwise alignments excluding the gap positions.

**Table S1. Amino acid sequence of purified Dsk2 constructs.** Color coding: UBL (blue), hotspot (HS) regions (orange), STI1 domain (red), and UBA (grey).

| Construct | Amino acid sequence |
|---|---|
| Dsk2 FL | MSLNIHIKSGQDKWEVNVAPESTVLQFKEAINKANGIPVANQRLIYSGKILKDDQTVESYHIQDGHSVHLVKSQPKPQTASAAGANNATATGAAAGTGATPNMSSGQSAGFNPLADLTSARYAGYLNMPSADMFGPDGGALNNDSNNQDELLRMMENPIFQSQMNEMLSNPQMLDFMIQSNPQLQAMGPQARQMLQSPMFRQMLTNPDMIRQSMQFARMMDPNAGMGSAGGAASAFPAPGGDAPEEGSNTNTTSSSNTGNNAGTNAGTNAGANTAANPFASLLNPALNPFANAGNAASTGMPAFDPALLASMFQPPVQASQAEDTRPPEERYEHQLRQLNDMGFFDFDRNVAALRRSGGSVQGALDSLLNGDV |
| Dsk2 I45A | MSLNIHIKSGQDKWEVNVAPESTVLQFKEAINKANGIPVANQRLAYSGKILKDDQTVESYHIQDGHSVHLVKSQPKPQTASAAGANNATATGAAAGTGATPNMSSGQSAGFNPLADLTSARYAGYLNMPSADMFGPDGGALNNDSNNQDELLRMMENPIFQSQMNEMLSNPQMLDFMIQSNPQLQAMGPQARQMLQSPMFRQMLTNPDMIRQSMQFARMMDPNAGMGSAGGAASAFPAPGGDAPEEGSNTNTTSSSNTGNNAGTNAGTNAGANTAANPFASLLNPALNPFANAGNAASTGMPAFDPALLASMFQPPVQASQAEDTRPPEERYEHQLRQLNDMGFFDFDRNVAALRRSGGSVQGALDSLLNGDV |
| Dsk2 ΔSTI1 | MSLNIHIKSGQDKWEVNVAPESTVLQFKEAINKANGIPVANQRLIYSGKILKDDQTVESYHIQDGHSVHLVKSQPKPQTASAAGANNATATGAAAGTGATPNMSSGQSAGFNPLADLTSARYAGYLNMPSADMFGPDGGALNNDAGMGSAGGAASAFPAPGGDAPEEGSNTNTTSSSNTGNNAGTNAGTNAGANTAANPFASLLNPALNPFANAGNAASTGMPAFDPALLASMFQPPVQASQAEDTRPPEERYEHQLRQLNDMGFFDFDRNVAALRRSGGSVQGALDSLLNGDV |
| Dsk2 ΔHS1 | MSLNIHIKSGQDKWEVNVAPESTVLQFKEAINKANGIPVANQRLIYSGKILKDDQTVESYHIQDGHSVHLVKSQPKPQTASAAGANNATATGAAAGTGATPNMSSGQSAGFNPGPDGGALNNDSNNQDELLRMMENPIFQSQMNEMLSNPQMLDFMIQSNPQLQAMGPQARQMLQSPMFRQMLTNPDMIRQSMQFARMMDPNAGMGSAGGAASAFPAPGGDAPEEGSNTNTTSSSNTGNNAGTNAGTNAGANTAANPFASLLNPALNPFANAGNAASTGMPAFDPALLASMFQPPVQASQAEDTRPPEERYEHQLRQLNDMGFFDFDRNVAALRRSGGSVQGALDSLLNGDV |
| Dsk2 ΔHS2 | MSLNIHIKSGQDKWEVNVAPESTVLQFKEAINKANGIPVANQRLIYSGKILKDDQTVESYHIQDGHSVHLVKSQPKPQTASAAGANNATATGAAAGTGATPNMSSGQSAGFNPLADLTSARYAGYLNMPSADMFGPDGGALNNDSNNQDELLRMMENPIFQSQMNEMLSNPQMLDFMIQSNPQLQAMGPQARQMLQSPMFRQMLTNPDMIRQSMQFARMMDPNAGMGSAGGAASAFPAPGGDAPEEGSNTNTTSSSNTGNNAGTNAGTNAGANTAANPNAGNAASTGMPAFDPALLASMFQPPVQASQAEDTRPPEERYEHQLRQLNDMGFFDFDRNVAALRRSGGSVQGALDSLLNGDV |
| Dsk2 ΔHS3 | MSLNIHIKSGQDKWEVNVAPESTVLQFKEAINKANGIPVANQRLIYSGKILKDDQTVESYHIQDGHSVHLVKSQPKPQTASAAGANNATATGAAAGTGATPNMSSGQSAGFNPLADLTSARYAGYLNMPSADMFGPDGGALNNDSNNQDELLRMMENPIFQSQMNEMLSNPQMLDFMIQSNPQLQAMGPQARQMLQSPMFRQMLTNPDMIRQSMQFARMMDPNAGMGSAGGAASAFPAPGGDAPEEGSNTNTTSSSNTGNNAGTNAGTNAGANTAANPFASLLNPALNPFANAGNAASTGMPQPPVQASQAEDTRPPEERYEHQLRQLNDMGFFDFDRNVAALRRSGGSVQGALDSLLNGDV |

**Table S2.** Summary of SAXS-derived Rg values for Dsk2 constructs from Guinier fitting (full SAXS parameters in Table S3).

| **Construct** | **Rg** (Å) |
|---|---|
| Dsk2 FL[a] | 36.7 ± 0.2[b], 37.6 ± 0.1[c] |
| Dsk2 ΔHS1 | 36.2 ± 0.2[b] |
| Dsk2 ΔHS2 | 38.5 ± 0.2[b] |
| Dsk2 ΔHS3 | 41.1 ± 0.2[c] |
| Dsk2 I45A | 39.1 ± 0.2[b] |

[a] Dsk2 FL is treated as an average $R_g$ of 37.2 ± 0.5 Å to account for variation from different beamline sessions

[b] and [c] represents SAXS datasets from different beamline sessions.

**Table S3. SAXS Data Collection Details**.

| *(a)* **Sample details** | | | | | | |
|---|---|---|---|---|---|---|
| Organism | *S. cerevisiae* | | | | | |
| Source (Catalogue No. or reference) | Expressed in *E. coli* (this work) | | | | | |
| Description: sequence (including Uniprot ID + uncleaved tags), bound ligands/modifications, etc. | Full-length Dsk2 (no tags), Uniprot ID P48510 | Dsk2 ΔHS1 (no tags) | Dsk2 ΔHS2 (no tags) | Dsk2 I45A (no tags) | Full-length Dsk2 (no tags), Uniprot ID P48510 | Dsk2 ΔHS3 (no tags) |
| Extinction coefficient ε in $M^{-1}$ $cm^{-1}$ (wavelength in nm) | 12950 (280) | 9970 (280) | 12950 (280) | 12950 (280) | 12950 (280) | 12950 (280) |
| Molecular mass M from chemical composition (Da) | 39,345 | 37,055 | 37,989 | 39,303 | 39,345 | 38,053 |
| For SEC-SAS, loading volume/concentration (mg $ml^{-1}$), injection volume (μl), flow rate (ml $min^{-1}$) | 6.11, 200, 0.6 | 6.49, 200, 0.6 | 6.32, 200, 0.6 | 6.11, 200, 0.6 | 6.11, 300, 0.6 | 6.35, 300, 0.6 |
| Solvent composition and source | pH 6.8 20 mM NaPhos | | | | | |
| *(b)* **SAS data collection parameters** | | | | | | |
| Instrument | BioCAT (Sector 18, APS), with Pilatus3 X 1M detector (Dectris) | | | | | |

| | | | | | | |
|---|---|---|---|---|---|---|
| Wavelength (Å) | 1.033 | 1.033 | 1.033 | 1.033 | 1.033 | 1.033 |
| Camera length (m) | 3.702 | 3.702 | 3.702 | 3.702 | 3.703 | 3.703 |
| *q*-measurement range | 0.0032-0.33 | 0.0032-0.33 | 0.0032-0.33 | 0.0032-0.33 | 0.0024-0.33 | 0.0024-0.33 |
| Normalization | Transmitted intensity | Transmitted intensity | Transmitted intensity | Transmitted intensity | Transmitted intensity | Transmitted intensity |
| Exposure time/number | 0.5 seconds (2376 frames) | 0.5 seconds (2470 frames) | 0.5 seconds (2470 frames) | 0.5 seconds (2470 frames) | 0.5 seconds (2376 frames) | 0.5 seconds (2376 frames) |
| Sample Configuration | SEC-MALS-SAXS using a Shodex KW-803 and an Agilent 1260 Series HPLC. UV data was measured with an Agilent 1290 DAD, and MALS/RI data by DAWN HELEOS-II (18-angle) and Optilab T-rEX (RI) instruments (Wyatt Technology). SAXS data was measured in a 1 mm, Mica-windowed flow cell | | | | | |
| Sample Temperature | 25 °C | | | | | |
| ***(c)* Software employed** | | | | | | |
| SAXS data reduction | Radial averaging; frame comparison, averaging, and subtraction done using BioXTAS RAW 2.4.0 or 2.4.1 [3] | | | | | |
| Basic analysis: Guinier, M.W. P(r) | Guinier fit and M.W. using BioXTAS RAW. RAW uses MoW and Vc M.W. methods [4,5] | | | | | |
| MALS-RI analysis | Astra 7 (Wyatt) | | | | | |
| ***(d)* Structural Parameters** | | | | | | |
| Guinier Analysis | Full-length Dsk2 | Dsk2 ΔHS1 | Dsk2 ΔHS2 | Dsk2 I45A | Full-length Dsk2 | Dsk2 ΔHS3 |

| | | | | | | |
|---|---|---|---|---|---|---|
| I(0) | 0.0157 | 0.0129 | 0.0154 | 0.0162 | 0.0212 | 0.0205 |
| $R_g$ (Å) | 36.7 ± 0.2 | 36.2 ± 0.2 | 38.5 ± 0.2 | 39.1 ± 0.2 | 37.6 ± 0.1 | 41.1 ± 0.2 |
| q-range ($Å^{-1}$) | 0.00494–0.02808 | 0.00663–0.02808 | 0.00381 – 0.02639 | 0.0055–0.02639 | 0.00663–0.02751 | 0.00353–0.02384 |
| Quality-of-fit parameter (with definition) | 0.9826 ($r^2$) | 0.9902 ($r^2$) | 0.9750 ($r^2$) | 0.9914 ($r^2$) | 0.9952 ($r^2$) | 0.9902 ($r^2$) |
| *M* from MALS (kDa) | 37 | 35 | 38 | 37 | 40 | 40 |

**Table S4.** Domain boundaries of the multidomain proteins used in this study. Brackets indicate the first and last residue of the domain.

| Uniprot code | Gene | N | Domain 1 | Domain 2 | Domain 3 | Domain 4 |
|---|---|---|---|---|---|---|
| P48510 | Yeast Dsk2 | 373 | [1,75] | [147,223] | [327,373] | |
| P48510 | Yeast Dsk2_ΔHS1 | 352 | [1,75] | [124, 202] | [306, 352] | |
| P48510 | Yeast Dsk2_ΔHS2 | 360 | [1,75] | [145, 223] | [314, 360] | |
| P48510 | Yeast Dsk2_ΔHS3 | 362 | [1,75] | [145, 223] | [316, 362] | |
| Q9UMX0 | Human UBQLN1 | 589 | [37,107] | [182,251] | [387,470] | [542,585] |
| Q9UHD9 | Human UBQLN2 | 624 | [33,103] | [178,247] | [379,462] | [577,620] |
| Q9NRR5 | Human UBQLN4 | 601 | [13,83] | [192,261] | [393,476] | [554,597] |
| Q9SII8 | Plant Dsk2B | 551 | [19,93] | [168,236] | [381,449] | [504, 548] |
| Q9SII9 | Plant Dsk2A | 538 | [18,93] | [163,231] | [364,444] | [491,535] |
| Q9JJP9 | Rat UBQLN1 | 582 | [28,102] | [173,245] | [381,457] | [539,579] |
| D4AA63 | Rat UBQLN2 | 638 | [30,103] | [190,263] | [393,469] | [592,635] |
| D4A3P1 | Rat UBQLN4 | 595 | [10,82] | [186,255] | [387,456] | [549,592] |
| Q9VWD9 | Fly UBQLN | 547 | [9,79] | [135,207] | [322,401] | [499,541] |
| G5EFF7 | C. Elegans UBQLN1 | 502 | [8,83] | [132,200] | [311,381] | [455,501] |
| Q8R317 | Mouse UBQLN1 | 582 | [26,100] | [174,242] | [382,454] | [535,579] |
| Q9QZM0 | Mouse UBQLN2 | 638 | [30,103] | [189,258] | [393,470] | [592,634] |
| Q99NB8 | Mouse UBQLN4 | 596 | [11,83] | [197,256] | [398,456] | [550,593] |
| Q4G000 | Zebrafish UBQLN | 599 | [25,97] | [186,255] | [398,456] | [552, 597] |
| F6RXL5 | Frog UBQLN1 | 564 | [17,89] | [165,233] | [372, 430] | [518,560] |
